# A Molecular and Cellular Mechanism for Bitter Taste in the mosquito *Aedes aegypti*

**DOI:** 10.64898/2026.06.28.734668

**Authors:** Leisl I. Brewster, Joshua U. Abel-Nwachukwu, Pei-Hsuan Wu, Oleksandr Hulai, Nicholas K. Tochor, Alyxzelle J. Relao, Cassidy S. Mark, Anjali Pandey, Trevor R. Sorrells, Benjamin J. Matthews

## Abstract

The yellow fever mosquito, *Aedes aegypti*, is a vector of Zika, Dengue, Chikungunya and yellow fever, disease-causing viruses impacting millions of people annually across the globe. Female mosquitoes transmit pathogens through serial blood-feeding, while both male and female mosquitoes feed on plant resources. Contact chemosensation (taste) guides these two feeding modes in both positive ways, feeding readily on sugar in nectar or ATP in blood, and negative ways, demonstrating aversion to chemically diverse bitter compounds in blood or sugar meals. Here, we identify a receptor, *AaegGr14*, that is expressed in neurons in the labellum and cibarium of the mosquito. Activation of *Gr14-*expressing neurons causes reduced feeding, while *Gr14* and *Gr14*-expressing GRNs are required for aversion to bitter compounds during nectar- but not blood-feeding. This work provides a molecular and cellular on-ramp to understand bitter taste in mosquitoes and establishes that there are feeding context-dependent differences in how taste mechanisms influence blood- and nectar-feeding. Understanding the molecular and cellular basis of bitter taste in mosquitoes during nectar- and blood-feeding will help inform vector control strategies and elucidate shared principles and unique aspects of insect taste systems.

## Introduction

Females of many mosquito species, including *Ae. aegypti*, require a blood-meal which they obtain from biting humans and other animals to develop their eggs (Clements, 1992). Plants are also an important food source for mosquitoes. Both sexes rely on plants to acquire essential nutrients and carbohydrates needed for flying, mating and host seeking (Barredo and DeGennaro, 2020; Foster, 1995; Peach and Gries, 2020). These include things such as nectar from flowers and extrafloral nectaries, fruits and plant tissue exudates and for males, this is the only meal that they consume (Clements, 1992). Mosquitoes are guided towards food sources using olfactory, visual and thermal cues (Coutinho-Abreu et al., 2022; Montell, 2025; Shannon et al., 2024). However, once they encounter the food source, they rely on their taste system to make the final decision about whether to accept or reject a meal (Baik and Carlson, 2020; Laursen, 2026). Given their distinct feeding modes (i.e. nectar- and blood-feeding), it is likely that mosquitoes deploy context-specific taste processing of positive and aversive taste cues.

Taste quality can provide information to animals about the chemical composition and potential value of potential food items. Sweet and umami taste, (and low concentrations of salt) generally promote feeding behaviour, while bitter and sour compounds (and high concentrations of salt) tend to supress feeding (Yarmolinsky et al., 2009). Bitter taste is important to prevent the ingestion of potentially toxic compounds (Glendinning, 1994; Yarmolinsky et al., 2009) including plant secondary metabolites such as alkaloids, terpenoids and phenolic compounds (Baker, 1977; Lüttge, 1977), which are likely produced as defence against predation by herbivores and insect pests (Adler, 2000; Biere et al., 2004; Johnson et al., 2006). While some of these bitter compounds simply repel insects once ingested, others have an insecticidal effect (reviewed in Ibanez et al., 2012) and insects have therefore evolved mechanisms to detect and avoid the ingestion of bitter chemicals (Bernays, 1998).

Mosquito taste (as in other insects) is distributed across multiple sense organs in mosquitoes, including tarsi, their mouthparts (notably the labellum and labrum), and an internal taste organ known as the cibarium (Baik and Carlson, 2020; Clements, 1992; Ignell and Hansson, 2005; Laursen, 2026; Mclver, 1982). Sensilla housing 3-5 gustatory receptor neurons (GRNs) are found on the surface of the external taste organs (Baik and Carlson, 2020; Laursen, 2026; McIver and Siemicki, 1978; Pappas and Larsen, 1976) and these GRNs project to higher order neurons in the ventral nerve cord (VNC; Matthews et al., 2019) or the suboesophageal zone (SEZ; Jové et al., 2020a).

Electrophysiological recordings from sensilla on the labellum of several mosquito species have provided insights into the cellular mechanisms of bitter detection and taste more broadly, showing that individual GRNs within a single sensillum respond to distinct taste modalities including bitters, sugars, salts and water (Baik et al., 2024; Kessler et al., 2013; Owen et al., 1974; Sanford et al., 2013; Sparks and Dickens, 2016a, 2016b). Individual bitter GRNs can respond to several bitter stimuli as well as other aversive chemicals such as the repellents DEET and Picardin (Sanford et al., 2013; Sparks and Dickens, 2016a, 2016b). Furthermore, adding bitter compounds to sugar or water could reduce neuronal responses in sugar or water sensitive GRNS (Baik et al., 2024; Kessler et al., 2013; Sparks and Dickens, 2016b).

The molecular mechanisms governing physiological responses within mosquito bitter GRNs remain unclear. To date, no bitter chemoreceptors have been identified in mosquitoes, and it is not clear how a single GRN is able to respond to a range of chemically diverse molecules. Research from *Drosophila melanogaster* suggests that insects may employ a complex and combinatorial code to encode bitter stimuli. Bitter compounds are detected by members of the Gustatory receptor (Gr) gene family, which comprise a diverse and early branching clade of chemoreceptor subunits (Clyne et al., 2000; Robertson, 2019; Robertson et al., 2003) forming homo- or hetero-tetrameric ligand-gated ion channels (Gomes et al., 2024; Ma et al., 2024). In *D. melanogaster*, multiple Grs are expressed in bitter sensitive gustatory receptor neuron, including a shared set of commonly expressed receptors (CERs) in all bitter GRNs as well as Grs unique to a given neuron (Dweck and Carlson, 2020; Weiss et al., 2011). The co-expression of different combinations of GRs produces a heterogeneous population of GRNs on the fly labellum, each of which respond to a wide range of bitter compounds (Delventhal and Carlson, 2016; Dweck and Carlson, 2020; Sung et al., 2017).

*Aedes aegypti* has a robust set of gustatory receptors with 72 annotated genes (Kent et al., 2008; Matthews et al., 2018). Orthology analysis reveals some conservation with *D. melanogaster*, notably in conserved and functionally validated receptors for carbon dioxide (McMeniman et al., 2014) and sugar (Jové et al., 2020a). However, there is also substantial divergence between Grs found in *D. melanogaster* and *Ae. aegypti*, as chemoreceptor gene families tend to evolve quickly and readily between species (Robertson, 2019). *D. melanogaster Gr66a* is a broadly expressed CER which labels many bitter neurons (Marella et al., 2006; Thorne et al., 2004; Wang et al., 2004) and is necessary for GRN responses to many bitter compounds (Dweck and Carlson, 2020; Lee et al., 2009; Moon et al., 2006; Shim et al., 2015). *Ae. aegypti* mosquitoes possess a one-to-one orthologue of *Gr66a* called *AaegGr14* (Matthews et al., 2018, 2016; Sparks et al., 2013), however, the role of this receptor in bitter taste has not been investigated. Here we generate and use a knock-in/knock-out allele of *Gr14* to identify the neurons expressing this receptor and determine its role in nectar and blood feeding. We show that *Gr14* contributes to bitter avoidance during nectar-feeding but is dispensable during blood feeding, revealing context-dependent mechanisms of bitter taste in mosquitoes.

## Results

### Gr14 is expressed in gustatory receptor neurons in the labellum and cibarium

To better understand the molecular and cellular basis of bitter taste in *Ae. aegypti* mosquitoes, we identified a clear 1:1 orthologue of *Drosophila melanogaster* Gr66a, *AaegGr14* (referred to hereafter as simply *Gr14*) (**Figure 1A**). We used CRISPR/Cas9-mediated mutagenesis to generate a knock-in/knock-out allele, *Gr14*^*QF2*^, which contains a QF2 transcriptional activator inserted into the coding region of exon 1 (Herre et al., 2022; Matthews et al., 2019; **Figure 1B**). When crossing *Gr14*^*QF2*^ to QUAS-dTomato-T2A-GCaMP6s, we identified expression within sensory neuron cell bodies within the labellum of female mouthparts (**Figures 1C**), as well as expression in male labellum and male and female cibarium (**Figure S1**). Within the brain, projections of *Gr14*-expressing cells terminated within the subesophegeal zone (SEZ), a primary taste center (**Figure 1D**). No expression was observed in tarsi or ventral nerve cord of male or female mosquitoes (data not shown). Together, the expression pattern of *AaegGr14*^*QF2*^ supports its role as a receptor involved in mosquito feeding behaviours.

**Figure 1:**
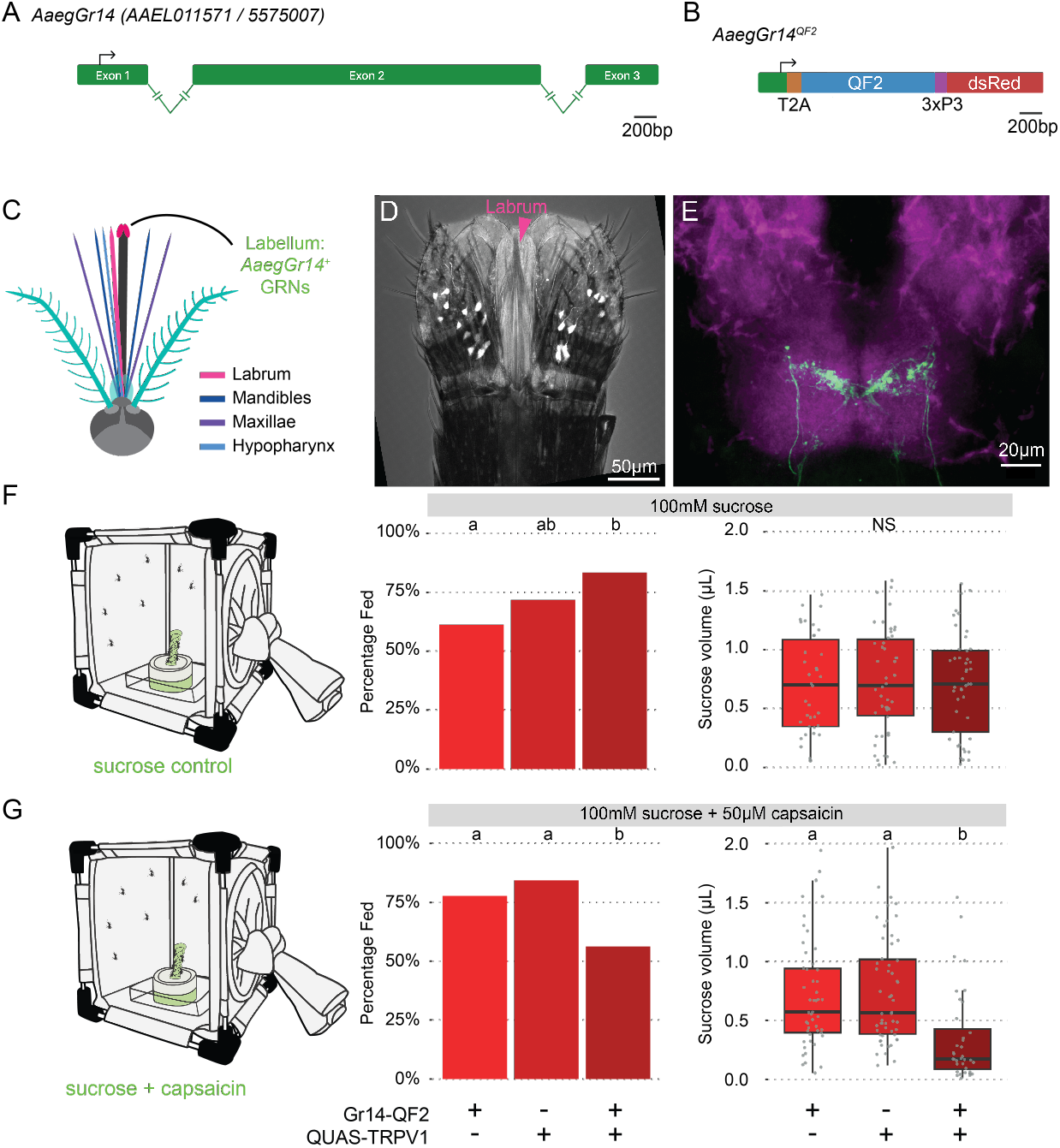
Generation and characterization of an *Aedes aegypti Gr14* knock-in/knock-out mutant and driver line. **(A)** Gene model of *AaegGr14* (alternative IDs: *AAEL011571* / 5575007). Introns are not to scale. **(B)** Resulting *AaegGr14*^*QF2*^allele generated by CRISPR/Cas9 homology-dependent repair. A T2A ‘ribosomal skipping peptide’ is in frame, and translation is predicted to result in a truncated AaegGr14 and a separate, full-length QF2 transcriptional activator. **(C)** Diagram of the mouthparts of *Ae. aegypti* mosquitoes. **(D)** Image of female labellum and labrum from *AaegQF2* crossed to *QUAS-dTomato-T2A-GCaMP6s* (Matthews et al., 2019) shows gustatory receptor neurons (GRNs) labelled with QF2-driven fluorescence. **(E)** Immunohistochemical localization of *Gr14*-expressing GRN axons in the subesophogeal zone (SEZ) of the female brain **(F)** Schematic of the single choice feeding assay where mosquitoes are offered a solution of 100mM sucrose, 0.1%DMSO and 0.02% fluorescein (Control) or a solution of 100mM sucrose +50µM capsaicin and 0.02% fluorescein (Capsaicin). **(G)** Percent-age of mosquitoes that ingested a control meal in 4 hours after a 24-hour water starvation. **(H)** Percentage of mosquitoes that ingested a capsaicin meal in 4 hours after a 24-hour water starvation. The genotypes included are driver controls (Gr14-QF2), reporter controls (QUAS-Kir2.1) and mosquitoes where Gr14 neurons are silenced (Gr4 > Kir2.1), 54-64 mosquitoes per genotype per treatment. **(I)** The volume of the control meal ingested by mosquitoes scored as fed. **(J)** The volume of the capsaicin meal ingested by mosquitoes scored as fed. Letters indicate significant differences between genotypes (in **(G-H)** logistic regression followed by Tukey HSD, in **(I-J)** Kruskall-Wallis test followed by Dunn’s test with a Bonferroni adjustment for multiple comparisons (*P* < 0.05).

### Activating GR14-expressiong neurons causes mosquitoes to reject a sucrose meal

To establish whether *Gr14-*expressing neurons have a role in mediating aversion during feeding, we chemogentically activated *Gr14* neurons by expressing the capsaicin sensitive receptor TRPV1 (Jové et al., 2020) in *Gr14* neurons. We found that adding 50µM capsaicin to a 10% sucrose meal significantly reduced the probability of mosquitoes consuming the meal when TRPV1 was expressed in *Gr14* neurons (**Figure 1E**). Furthermore, when mosquitoes did consume a meal, they ingested a significantly lower volume when *Gr14* neurons were activated by capsaicin compared to genetic controls (**Figure 1F**). Taken together, these results show that activation of *Gr14* neurons is sufficient to cause aversion to an otherwise appetitive stimulus.

### Gr14 mediates detection of bitters during nectar-feeding

To determine whether *Gr14* receptors are involved in the detection of bitter substances in a nectar-feeding context, we tested whether *Gr14*^*QF2/QF2*^ mutant mosquitoes were impaired in their ability to detect bitter substances. We presented mosquitoes with a choice between a sucrose only control and a stimulus containing sucrose at the same concentration mixed with bitter compounds of varying concentrations (**Figure 2A-B**). The choices of bitter compound and concentrations were informed by previous studies in *Ae. aegypti* (Ignell et al., 2010), *Aedes albopictus* (Baik et al., 2024), and *D. melanogaster* (Weiss et al., 2011). We confirmed that dye colour did not affect preference by presenting two sucrose stimuli as a control **(Figure S2**). The genotypes showed no difference in preference when presented with sucrose alone however, wildtype and driver control mosquitoes showed aversion to escin, lobeline, denatonium and quinine at the indicated concentrations (**Figure 2C**). *Gr14*^*QF2/QF2*^ mutant mosquitoes showed significantly reduced aversion to escin and quinine as compared to wild-type or heterozygous *Gr14*^*QF2/+*^ mosquitoes. We also observed a reduction in aversion to lobeline in *Gr14* mutants, however, they still maintained aversion to this compound. While we observed a trend of reduced aversion of *Gr14*^*QF2/QF2*^ mosquitoes to denatonium this was not statistically significant. Caffeine presented at 10mM did not significantly impact feeding preference in any of the genotypes (**Figure 2C**). Across all treatments the feeding rates of *Gr14* ^*QF2/QF2*^ mosquitoes were similar to those of controls (**Figure S3**).

**Figure 2.**
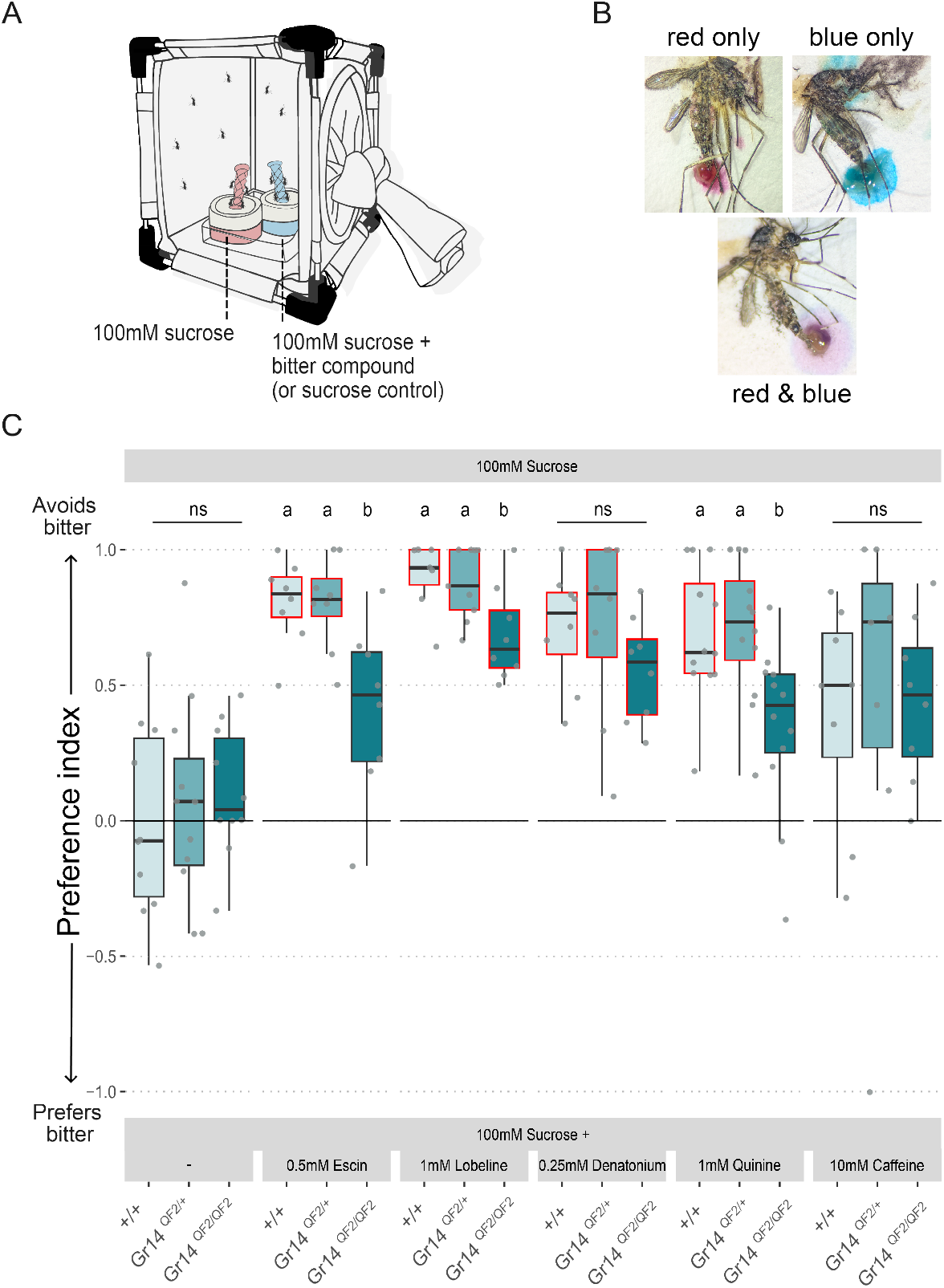
*Gr14* is necessary for the avoidance of bitter substances during nectar feeding. **(A)** Schematic of the two-choice sucrose feeding assay, ∼15 non-blood-fed females/cage **(B)** Females who have ingested either the red, blue or a mix of both stimuli. **(C)** The feeding preferences of wildtype (+/+), heterozygotes (*Gr14*^*QF2/+*^) and homoallelic Gr14 mutants (*Gr14*^*QF2/QF2*^) for the indicated bitter substance (n = 7-12 trials). Feeding preference is indicated by a preference index, where 1.0 = complete preference for sucrose and −1.0 = a complete preference for a mix of sucrose/stimulus. Boxplots are showing the median and upper and lower quartiles. Letters above each box (top row) indicate significant PI differences between genotypes (Kruskall-Wallis test followed by Dunn’s test with a Bonferroni adjustment for multiple comparisons (*P* < 0.05)). Red boxes indicate significance between sucrose controls and a given bitter treatment across genotypes (Kruskall-Wallis test followed by Dunn’s test with a Bonferroni adjustment for multiple comparisons).

### Silencing *Gr14*-expressing neurons has different effects with different bitter compounds

Next, we wanted to examine the role of *Gr14*-expressing neurons in mediating the detection of bitter substances. We silenced *Gr14*-expressing neurons by expressing the inwardly rectifying potassium channel Kir2.1. We generated and tested a QUAS-Kir2.1 effector line by first crossing it to a Gr4-QF2 driver, which labels putative sweet-detecting GRNs in the labellum of the mosquito (Jové et al., 2020a). Silencing Gr4 neurons reduces sugar intake **(Figure S4**), indicating that this QUAS-Kir2.1 line is effective in silencing neural activity.

We crossed QUAS-Kir2.1 to our Gr14 driver and examined feeding preference using our two-choice assay with sucrose and sucrose laced with bitter compounds. Silencing *Gr14-*expressing neurons had no effect on sucrose preference in the sucrose only controls, while silencing *Gr14*-expressing neurons reduced aversion to escin and denatonium to levels indistinguishable from control (**Figure 3A**). Silencing *Gr14-*expressing neurons also reduced aversion to lobeline, however, when comparing this aversion to sucrose only controls a significant aversion remained. Silencing Gr14 neurons did not affect quinine or caffeine preference. Across all treatments, silencing Gr14 neurons did not affect the feeding rate (**Figure S3**).

**Figure 3.**
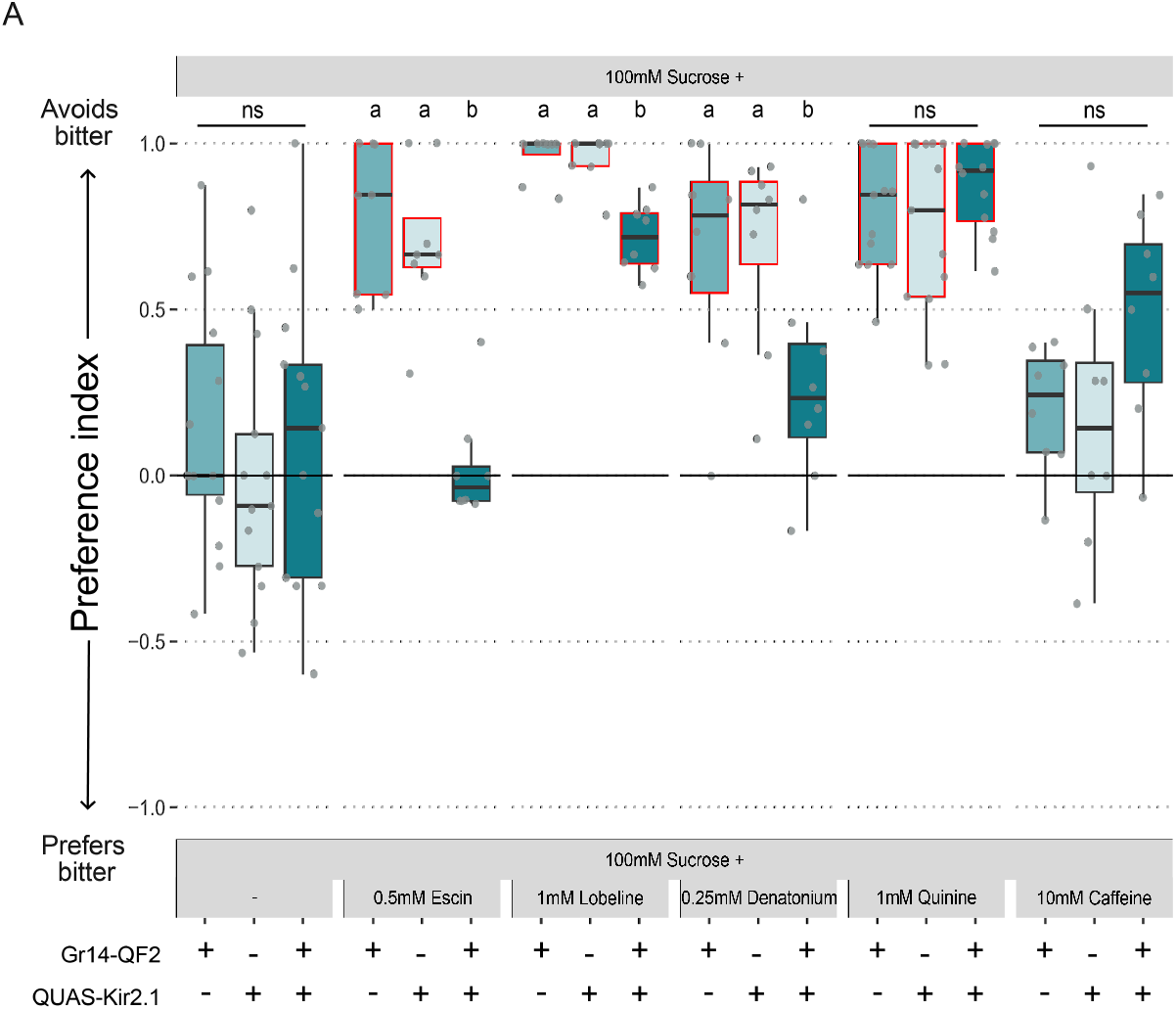
*Gr14*-expressing neurons are necessary for the avoidance of some bitter substances in a sugar feeding context. **(A)** The feeding preferences driver controls (*Gr14-QF2*), reporter controls (*QUAS-Kir2*.*1*) and mosquitoes where *Gr14* expressing neurons are silenced (*Gr14 > Kir2*.*1*) for the indicated bitter substance (n = 8-12 trials). Feeding preference is indicated by a preference index where 1.0 = complete preference for sucrose and −1.0 = a complete preference for a mix of sucrose/stimulus. Boxplots are showing the median and upper and lower quartiles. Letters above each box (upper row) indicate significant PI differences between genotypes (Kruskall-Wallis test followed by Dunn’s test with a Bonferroni adjustment for multiple comparisons (*P* < 0.05)). Red boxes indicate significance between sucrose controls and a given bitter treatment across genotypes (Kruskall-Wallis test followed by Dunn’s test with a Bonferroni adjustment for multiple comparisons).

### Detection of bitter compounds directly in blood meals is independent of *Gr14*

To test whether *Gr14* receptors are required for aversion to bitter compounds during blood feeding, we first performed a two-choice In-Meal assay in which mosquitoes were offered either a control substitute blood meal (minimal meal) or a minimal meal supplemented with a bitter tastant **(Figure 4A-C)**. When given a choice between two control meals, neither genotype displayed a feeding preference. However, the addition of 1mM escin or lobeline, as well as 10 mM denatonium, significantly shifted feeding preference toward the minimal meal-only stimulus in both genotypes. In contrast, neither 1mM of caffeine nor quinine strongly deterred feeding under these conditions **(Figure 4D)**. Across all compounds tested, *Gr14* mutants showed aversion comparable to wildtype mosquitos **(Figure 4D)**. Consistent with this, 1 mM of quinine similarly reduced minimal meal intake in a single-choice feeding assay in both genotypes, while consumption of a control minimal meal did not differ between genotypes **(Figure S5)**.

**Figure 4.**
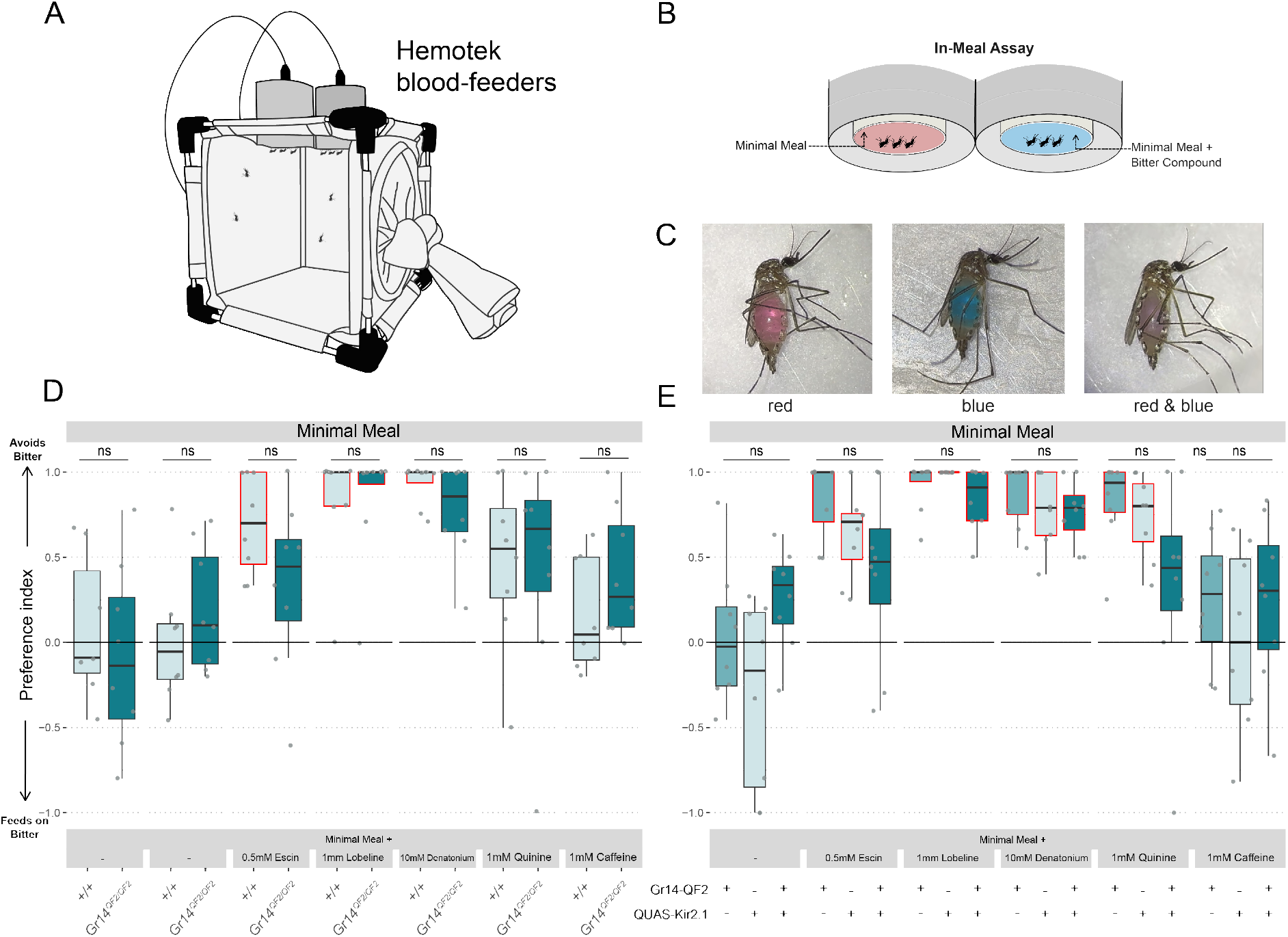
Detection of bitter compounds directly in blood meals does not require *Gr14*. **(A-B)** Schematic of two-choice in-meal feeding assay; mosquitoes choose between a minimal meal or a minimal meal supplemented with a given bitter compound. **(C)** Scoring of mosquitos who have ingested either the red, blue or a mix of both stimuli. **(D)** The feeding preferences of wildtype (+/+),and homoallelic *Gr14* mutants (*Gr14*^*QF2/QF2*^) for the indicated bitter substance (n = 8 trials of 9-12 mosquitos per cage). **(E)** The feeding preferences driver controls (*Gr14-QF2*), reporter controls (*QUAS-Kir2*.*1*) and mosquitoes where *Gr14* expressing neurons are silenced (*Gr14 > Kir2*.*1*) for the indicated bitter substance (n = 8 trials of 9-12 mosquitos per cage). Feeding preference is indicated by a preference index where 1.0 = complete preference for minimal meal and −1.0 = a complete preference for a mix of minimal meal and bitter stimulus. Boxplots are showing the median and upper and lower quartiles. Letters above each box indicate significant PI differences between genotypes (Kruskall-Wallis test followed by Dunn’s test for multiple comparison (*P* < 0.05)). Red boxes indicate significance between sucrose controls and a given bitter treatment across genotypes (Kruskall-Wallis test followed by Dunn’s test with a Bonferroni adjustment for multiple comparisons).

We next asked whether silencing *Gr14*-expressing neurons impairs the detection of bitter compounds directly in blood meals. In driver control mosquitoes, 0.5mM of escin, 1mM of lobeline, 10mM of denatonium, and 1mM quinine all significantly shifted feeding preference toward the minimal meal only stimulus. Effector controls showed a similar pattern of avoidance, though with escin no longer significantly affecting preference. In mosquitoes with Gr14 neurons silenced, only lobeline and denatonium significantly altered feeding preference relative to controls **(Figure 4E)**. Across bitter compounds, silencing *Gr14*-neurons did not significantly alter avoidance of bitter compounds directly in blood meals **(Figure 4E)**.

### Surface detection of bitter compounds during blood-feeding does not require *Gr14*

Next, we asked whether *Gr14* receptors are required for aversion to bitter compounds encountered on feeding surfaces. We modified the two-choice assay to a Surface Contact assay, in which mosquitoes chose between identical minimal meals delivered through a Kimwipe coated with either water or a bitter tastant **(Figure 5A-B)**. In wildtype mosquitoes, 100 mM of lobeline and quinine, as well as 10 mM of escin significantly shifted feeding preference toward the water-only stimulus. 100mM of Denatonium and caffeine did not significantly alter preference. In *Gr14* mutants, only 100 mM of lobeline produced an aversive effect. However, when comparing genotypes, preference indices did not significantly differ for any bitter compound **(Figure 5C)**.

**Figure 5.**
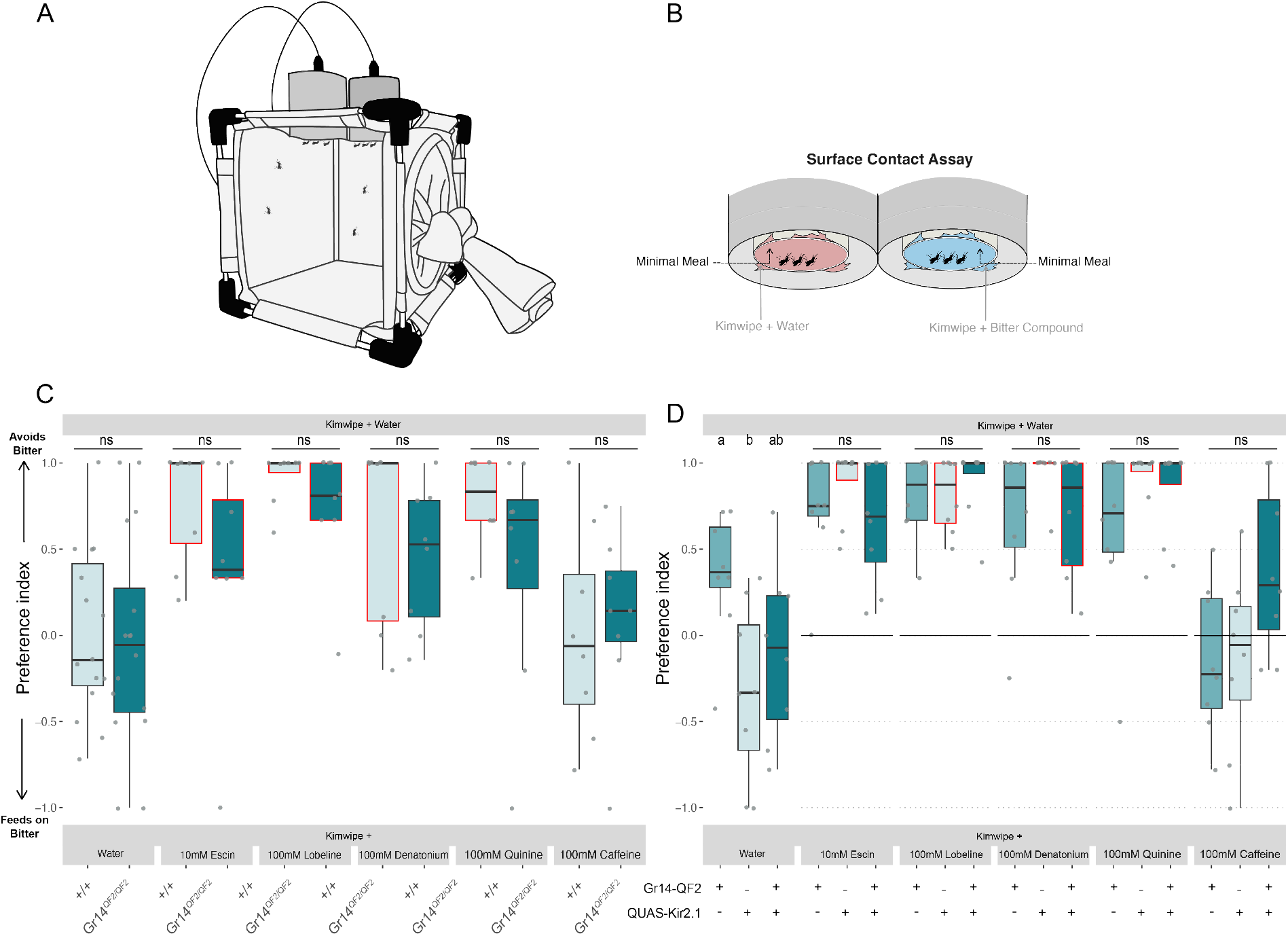
Detection of bitter compounds on the surface of a blood meal does not require *Gr14*. **(A)** Schematic of two-choice Surface Contact assay; Mosquitoes contact a Kimwipe coated with water or with an indicated bitter compound prior to feeding on control minimal meals. **(B)** The feeding preferences of wildtype (+/+) and homoallelic Gr14 mutants (*Gr14*^*QF2/QF2*^) for the indicated bitter substance (n = 8 trials of 9-12 mosquitos per cage). **(C)** The feeding preferences driver controls(*Gr14-QF2*),reporter controls (*QUAS-Kir2*.*1*) and mosquitoes where *Gr14* expressing neurons are silenced (*Gr14 > Kir2*.*1*) for the indicated bitter substance (n = 8 trials of 9-12 mosquitos per cage).Feeding preference is indicated by a preference index where 1.0 = complete preference for water on the surface and −1.0 = a complete preference for the bitter stimulus on the surface. Box-plots are showing the median and upper and lower quartiles. Letters above each box indicate significant PI differences between genotypes (Kruskall-Wallis test followed by Dunn’s test for multiple comparison (*P* < 0.05)). Red boxes indicate significance between sucrose control and a given bitter treatment across genotypes (Kruskall-Wallis test followed by Dunn’s test with a Bonferroni adjustment for multiple comparisons).

Finally, we tested whether silencing *Gr14*-expressing neurons affects surface-mediated detection of bitter compounds. During surface contact assays, addition of 100 mM lobeline, denatonium, or quinine, as well as 10 mM escin, significantly shifted feeding toward the water-coated Kim-wipe in effector controls and Gr14-silenced mosquitoes, but not in driver controls **(Figure 5D)**. Overall, silencing *Gr14* neurons did not significantly change feeding preference indices across bitter compounds **(Figure 5D**). Feeding rate differences between genotypes and tastants were minimal and did not correlate with particular genotypes or compounds (**Figure S6**).

## Discussion

Here, we demonstrate that mosquitoes possess redundant bitter taste pathways and process bitter cues differently depending on feeding context (nectar- vs. blood-feeding). We identify Gr14 as the first functionally characterized mosquito bitter receptor and show that it contributes to bitter avoidance during nectar-feeding, but is dispensable during blood feeding, thus revealing context-dependent mechanisms of gustatory processing. *Gr14* is expressed in neurons in the labellum and cibarium that can prevent feeding and that *Gr14* and *Gr14-*expressing neurons are necessary for a complete avoidance of several bitter compounds in nectar-feeding but not blood-feeding.

### *Gr14* mediates aversion to several bitter compounds

We found that *Gr14* was involved in mediating the aversion to several bitter compounds, namely escin, lobeline, denatonium benzoate and quinine in a nectar-feeding context. What is the mechanism by which a single receptor can mediate the detection of several different compounds? First, it is possible that Gr14 ligand binding domains can bind promiscuously to structurally diverse compounds. However, we find this possibility unlikely, given that bitter chemicals vary enormously in their molecular diversity (Ziaikin et al., 2025). Instead, we propose that *Gr14* may act as a co-receptor within heterometric receptor complex(es). In *D. melanogaster*, bitter detection is mediated by neurons that express a core set of commonly expressed receptors (CERs) and up to dozens of Grs unique to a given neuron (Dweck and Carlson, 2020; Weiss et al., 2011). These Grs likely assemble into heterotetrameric receptors with distinct tuning properties based on their subunit (Dweck and Carlson, 2020; Lee et al., 2009; Shim et al., 2015; Sung et al., 2017). We thus speculate that Gr14 can form multiple heteromeric receptor complexes with other Grs. Supporting this model, a snRNA-seq dataset suggests that *Gr14* is co-expressed with several other Gr sub-units in proboscis GRNs, providing potential partners for distinct hetero-tetrameric bitter receptors (Goldman et al., 2025).

### Redundant mechanisms in bitter detection

Our results suggest that mosquitoes employ redundant mechanisms for bitter detection. Such redundancy could be occurring through several (not mutually exclusive) mechanisms.

Firstly, additional bitter receptors within *Gr14-*expressing neurons that can compensate for the loss of Gr14. This may explain why in nectar-feeding experiments, knocking out *Gr14* did not significantly impact aversion to denatonium while silencing these neurons did. We also observed a similar trend in escin where silencing *Gr14* neurons seemed to result in a greater loss of aversion than in the mutant. In flies, there are examples where the loss of a bitter receptor was either partially (Dweck and Carlson, 2020) or completely compensated by the presence of another receptor (Sung et al., 2017). The ability to form multiple heteromeric complexes within a single GRN would facilitate such redundancy.

Secondly, non-*Gr14*-expressing bitter neurons may be compensating for the loss of these neurons. In the proboscis, Gr14 neurons are a subset of a broader GRN population characterized by the expression of *Gr20* (Goldman et al., 2025). Unlike *Gr14, Gr20* is expressed in the tarsi across a population of putative nociceptive neurons. These neurons also express *TRPA1*, a receptor that detects high temperatures and nociceptive chemicals (Corfas and Vosshall, 2015; Kwon et al., 2010; Melo et al., 2021). Aversion after silencing *Gr14-*expressing neurons may be maintained by non-*Gr14*-expressing neurons such as the *Gr20* and *TRPA1* expressing neurons in the tarsi, or by non-*Gr14-*expressing bitter neurons in the proboscis or cibarium.

Finally, aversion may be occurring through the suppression of neural responses within sugar sensitive GRNs. In *D. melanogaster*, addition of some bitter compounds to sugar reduces the neural activity in sugar sensitive neurons (French et al., 2015; Jeong et al., 2013). Suppression of sugar neuron activity by bitter compounds is also observed in *Aedes albopictus* (Baik et al., 2024) and *Anopheles gambiae* (Kessler et al., 2013) making it a plausible mechanism for bitter aversion in mosquitoes. Electrophysiological recordings in wild-type and *Gr14* mutant mosquitoes will be necessary to resolve the degree to which this inhibition contributes to bitter-mediated anti-feeding behaviour in mosquitoes.

### Aversion during blood-feeding is mediated primarily through non-*Gr14* mechanisms

While mosquitoes are more likely to encounter bitters in nectar-feeding, the role of bitters in blood-feeding has been investigated due to their potential as contact repellents added to the skin or as anti-feedants where aversive compounds are added to an artificial feeder (Baik et al., 2024; Dennis et al., 2019; Kessler et al., 2013; Lazzari et al., 2024). These studies found that bitter compounds added to blood or artificial blood-meals effectively reduce ingestion, however, substantially higher concentrations of bitter compounds are needed to prevent biting or blood-feeding when bitter compounds are added to the skin. Similarly, we found that a substantially higher concentration of the same bitter chemical was required to elicit aversion on the surface compared to when it was added to the blood-meal or when added to sugar.

Unlike in nectar-feeding, *Gr14* and *Gr14*-expressing neurons were not necessary for the aversion to bitter compounds during blood-feeding. This cannot be explained simply by lack of receptor contact with the substrate since across all three feeding assays, *Gr14*-expressing neurons are likely directly contacting bitter compounds using a variety of tissues: the cibarium and labellum during nectar-feeding, the cibarium during in-meal blood-feeding, and the labellum during surface contact blood-feeding. Instead, we propose that changes in feeding context may change which sensory input the mosquito prioritizes. The mosquito mouthparts are specialized for different modes of feeding, allowing them to differentiate between blood and sugar in the periphery. Sensory neurons in the labrum respond specifically to the different components of blood while sweet sensitive neurons in the labellum motivate nectar-feeding, and each population of neurons projects to anatomically distinct regions of the SEZ (Jové et al., 2020a). Thus, it is possible that tissues in which Gr14 is not expressed, such as the tarsi and the labrum, may be more salient during blood-feeding.

### Limitations and future directions

A broader range of concentrations of bitters in our panel as well as a larger panel of bitters will be necessary to determine its full role in bitter detection and aversion. Gr20 driver lines would be important. Additionally, electrophysiology or calcium imaging will be necessary to prove that *Gr14* is a redundant receptor. We propose a model where Gr14 (and likely Gr20) act as a co-receptor that assemble into heterotetrameric receptor complexes with other Gr subunits. Electrophysiological recordings from several sensilla on the proboscis when *Gr14* is mutated out or when *Gr14*-expressing neurons are silenced would test this hypothesis as well as having access to Gr20 mutants/QF2 lines. There may be strong evolutionary pressure to evolve such redundancy in bitter perception as the ingestion of bitters can result in large fitness deficits such as reduced egg output (Kessler et al., 2013) and death (Muñoz et al., 2020). More broadly, our findings suggest that mosquito bitter taste is organized not as a fixed sensory pathway, but as a context-dependent system in which distinct gustatory circuits are recruited during nectar and blood feeding.

## Supporting information

Raw data for Brewster et al. 2026

## Acknowledgements

The authors wish to thank members of the Matthews lab for feedback on this project and manuscript. We would like to acknowledge Meg Younger for the advice on immunostaining and imaging mosquito CNS. This work was supported by grants from Natural Sciences and Engineering Research Council of Canada (Discovery Grant RGPIN-2020-05423 to B.J.M.), Canadian Institutes of Health Research (Project Grant PJT-191985 to B.J.M.), Michael Smith Health Research BC (Scholar Award SCH-2021-1860 to B.J.M.), Alfred P. Sloan Foundation (Research Fellowship in Neuroscience FG-2021-16383 to B.J.M.), and Human Frontier Science Program (Research Grant RGP018/2023 to B.J.M. and a S4S supplement to O.H.). Additional infrastructure support was provided by the Canadian Foundation for Innovation and British Columbia Knowledge Development Fund (CFI JELF and BCKDF awards 40036, 41223, 43348 to B.J.M.) and unrestricted startup funds from The University of British Columbia (B.J.M.). T.S. is funded by NIH grant DP2AI177891. T.S. is an HHMI Freeman Hrabowski Scholar. This paper was typeset with the bioRxiv word template by @Chrelli: www.github.com/chrelli/bioRxiv-word-template

## Author contributions

L.I.B. designed research, performed sugar-feeding experiments, curated and analyzed data, generated figures, wrote the manuscript and led the project

J.U.A. designed research performed blood-feeding studies, analyzed data, generated figures, and contributed to writing and editing the manuscript.

L.I.B., C.M., O.H., and N.K.T. generated the Gr14^QF2^ mutant allele

O.H. contributed to genetic crosses and strain maintenance

P-H.W. performed all anatomical studies

A.P. and T.R.S. generated, tested, and mapped the QUAS-Kir2.1 line

A.J.R. and L.I.B. validated the QUAS-Kir2.1 in Gr4-expressing neurons

B.J.M. designed research, supervised the project, obtained funding, analyzed data, generated figures and contributed to writing and editing of the manuscript

All authors approved of the submission of this manuscript

## Competing interest statement

The authors declare no competing financial interests

## Materials and Methods

### Mosquito Rearing and Maintenance

All *Aedes aegypti* wild-type laboratory strains (Liverpool) and genetically modified strains were maintained and reared at 25–28°C, 70–80% relative humidity with a photoperiod of 14 hr light: 10 hr dark as previously described in (DeGennaro et al., 2013). Eggs were hatched in dechlorinated water under vacuum pump and larvae were kept in dechlorinated water and fed on a diet of fish food mixed with agar. Adult mosquitoes were provided constant access to 10% sucrose (w/v in dechlorinated water). For routine strain maintenance animals were blood-fed on a Hemotek membrane feeding system using defibrinated sheep’s blood heated to 37°C and eggs were collected on germination paper placed in dechlorinated water.

Female *Aedes aegypti* mosquitoes (Liverpool strain) used in these experiments were between 7-14 days old (except for the capsaicin feeding experiments; 2-14 days old) and were not blood fed unless otherwise specified by the experimental protocol. Females were housed in mixed sexed cages prior to experimental trials.

### Generation of Genetically Modified Mosquito Strains

The GR14 knock-in/knock-out strain was generated using CRISPR-Cas9 mediated homology-directed repair as previously described in (Kistler et al., 2015; Matthews et al., 2019). A guide RNA was designed to target exon 1 of the *AaegGr14* locus (target sequence with PAM underlined: 5’-GAC-GGATGCCTACGTCGGAG**TGG**-3’). sgRNA was prepared via *in vitro* transcription using the NEB EnGen sgRNA Synthesis kit, *S. pyogenes* (NEB E3322V), following the manufacturer’s protocol.

The donor plasmid was constructed using NEBuilder HiFi DNA Assembly (NEB E5520S) using the following fragments: homology arms of 1 kb on either side of the Cas9 cut site, a fragment containing *T2A-QF2-SV40* and *3xP3-dsRed*, PCR-amplified from a vector derived from ppk301-T2A-QF2 HDR plasmid (Addgene #130667; Matthews et al., 2019) and a *pUC19* backbone digested with BamHI and XBal restriction enzymes. Colonies were screened for transformation with the donor plasmid using colony PCR and the donor plasmid was purified using an endotoxin-free midiprep kit (Macherey Nagel 740422.50).

A total of 2067 Exu-Cas9 (Li et al., 2017) embryos were injected with a mixture of 80ng/ul sgRNA and 500ng/ul donor plasmid under aqueous media (Harrell, 2024). A total of 209 larvae hatched and were out-crossed to wild-type mosquitoes to generate F1 families to screen for germ-line integration of our cassette. Of 86 families that screened for the expression of dsRed in the optic nerve, four F1 families positive for 3xP3-dsRED expression were isolated and a complete insertion was verified using PCR followed by Sanger sequencing. One of these families was chosen and outcrossed for four generations to wild-type mosquitoes. The fourth generation was then crossed together for three generations, selecting mosquitoes with the most intense eye marker fluorescence to enrich for homozygosity. The genotype of the final homozygous mutant strain was verified by PCR.

Mosquitoes expressing the human Kir2.1 sequence under control of the QUAS promoter were created by cloning this sequence into the pXL-BacII backbone and integrating into the mosquito genome using the piggyBac transposase. The GFP-hKir2.1 sequence was amplified from UAS-FRT-CD2-FRT-Kir2.1-GFP from (Yang et al., 2009) using the primers 5’-CTCGAGCAAAATGGTGAGCAAGGGCGAGGAGCTG-3’ and 5’-ATCCTCTAGATCATATCTCCGATTCTCGCCGTAAGGGCC-3’. This insert was cloned into pXL-BacII-15xQUAS_TATA-SV40 (Riabinina et al., 2016) backbone, amplified using the primers 5’-TGCTCACCATTTT-GCTCGAGCCGCGGCCGCAGATC-3’ and 5’-GGAGATATGATCTAGAG-GATCTTTGTGAAGGAACCTTACTTCTG-3’. These components were combined with Infusion HD cloning kit (Takara 638920) to create pTS28 (QUAS-EGFP-hKir2.1). The plasmid was injected into 500 Liverpool strain *A. aegypti* embryos at the Insect Transformation Facility (Rockville, MD) using 200 ng/µL pTS28 plasmid and 200 ng/µL piggyBac transposase mRNA. Ten independent insertions were isolated under standard mosquito rearing conditions. Lines were crossed to a driver that expresses broadly in the nervous system and assessed for lethality to screen for efficacy. Lines were mapped in the genome using TagMapping (Stern, 2016). The QUAS-Kir2.1 line used in this paper (#8) mapped to chromosome 3.

### Immunohistochemistry

The immunostaining procedure was adapted from a previously published protocol (Younger, 2024). Adult mosquitoes of both sexes, 6 to 12 days post-emergence, were anesthetized on ice, after which they were carefully de-capitated using fine-tip spring scissors. The thoracic segment of the remaining body was also cut, and the legs and wings were removed. The heads and thoracic segments were immersed in a fixative solution containing 4% paraformaldehyde, 0.1 M Millonig’s Phosphate Buffer (pH 7.4), and 0.25% Triton X-100 for 3 hours at 4°C. The Millonig’s Phosphate Buffer was home-made following the recipe of Millonig (1964). The fixative was always freshly prepared, kept cold, and used within a week. After fixation, tissues were transferred into ice-cold 1× phosphate-buffered saline (PBS), diluted from 10× PBS (P5493, Sigma), which is free of Ca^2+^ and Mg^2+^. The 1× PBS has a phosphate buffer concentration of 0.01 M, a sodium chloride concentration of 0.154 M, and a pH of 7.4.

### Two Choice Sucrose Preference assay

The female mosquitoes used in all behavioural assays were mated and between 2 – 14 days old. Mosquitoes were starved of a sugar meal with access only to dechlorinated water for ∼24hrs prior to experimentation and ∼15 individuals were then transferred to small mesh cages (W17.5 x D17.5 x H17.5 cm – Bugdorm BD4M1515) where two choices of meals were presented. Mosquitoes were given a choice between 100mM sucrose solution and a mix of 100mM sucrose with given bitter tastant at a particular concentration and meals were either dyed red with 0.125% Amaranth (Sigma - Aldrich #A1016) or blue with 0.125% FC and C Blue No.1 (Spectrum #FD110). To control for dye preference the colour of the meals was swapped for half of the replicates. Mosquitoes were allowed to feed for 24 hours under standard rearing conditions (25–28°C, 70–80% relative humidity with a photoperiod of 14 hr light: 10 hr dark). Mosquitoes were frozen and a dissecting microscope was used to score the colour of the meals consumed by squashing the abdomen on a piece of white filter paper with a spatula.

### Minimal Meals

Artificial blood meals containing the minimum requirements to produce feeding responses similar to blood (minimal meals; Jové et al., 2020b) were used in blood feeding assays. For all assays, 10mL minimal meals containing a final concentration of 1mM ATP and 120mM NaHCO_3_ were prepared. Stimuli of interest and/or food or fluorescent dyes were added as necessary. 0.125% of either Amaranth Red or FD&C blue were used as food dyes for qualitative two choice assays, and 0.002% fluorescein was used for quantitative single-choice assays.

### Two Choice Blood In-Meal assays

Cages of mated, 7-14 day old female mosquitoes were starved on dechlorinated water for ∼24 hours prior to the assay. Mosquitos were presented with a control minimal meal or a minimal meal containing a bitter tastant, wrapped in parafilm, and allowed to feed for at least 15 minutes using a Hemotek feeder. Meal choice was evaluated via scoring abdomen color by eye. In cases where abdomen color was not obvious, mosquito abdomen was gently squished on a kimwipe to reveal dye. Meal Preference was assessed by calculating the number of mosquitos that chose the control meal, bitter meal, or both using PI = (Control-Bitter)/(Control+Bitter+Both).

### Two Choice Blood Surface Contact Assays

Cages of mated, 7–14-day old female mosquitoes were starved on dechlorinated water for ∼24 hours prior to the assay. Mosquitos were presented a minimal meal wrapped in a parafilm, and either a Kimwipe coated with water (control) or a Kimwipe coated with a bitter tastant, and allowed to feed for at least 30 minutes. Meal choice was evaluated via scoring abdomen color by eye. In cases where abdomen color was not obvious, mosquito abdomen was gently squished on a filter paper to reveal dye color. For coating of Kimwipe, 150µL of stimulus was applied to the Kimwipe for adhesion to the parafilm surface, then right before addition to the Hemotek another 150µL coating was applied to the Kimwipe to maintain moisture throughout the duration of feeding. Meal Preference was assessed by calculating the number of mosquitos that chose the control meal, bitter meal, or both using PI = (Control-Bitter)/(Control+Bitter+Both).

### Quantitative single choice blood-feeding assays

Fluorescein at a final concentration of 0.002% was added to 10mL of control minimal meal, or a minimal meal containing 1mM of quinine wrapped in parafilm and mosquitoes were allowed to feed for at least 1 hour using a Hemotek feeder.

### Single choice nectar-feeding assays

Mixed sex cages of 2-14 day old mosquitoes were starved of a sugar meal with access only to dechlorinated water for ∼24hrs prior to experimentation. Females were transferred to small mesh cages (W17.5 x D17.5 x H17.5 cm – Bugdorm BD4M1515) with access to 10% sucrose mixed with fluorescein at a final concentration of 0.002%. In experiments with capsaicin, mosquitoes were offered either a control meal consisting of 10% sucrose, 0.002% fluorescein and 0.01% DMSO (which was used as a vehicle to for capsaicin) or a meal consisting of 10% sucrose, 0.002% fluorescein and 50µM capsaicin (Tocris Bioscience - 0462/100). 32 females were not offered a fluorescein meal and reserved as unfed controls. Mosquitoes were allowed to feed for 4 hours. At the end of each trial mosquitoes were frozen at −20 °C until meal quantification was done.

### Single choice meal size quantification

Meal size quantification was done according to (Venkataraman et al., 2022). Animals that were offered a fluorescein meal as well as 8-16 unfed controls were added to a 96-well plate with one 2mm diameter stainless steel bead and 100µl of 1x PBS. 8 animals that were not offered a meal were used to generate a standard curve by adding the following volumes of 10% sucrose + 0.002% fluorescein: 5, 2.5, 1.25, 0.625, 0.3125, 0.15625, 0.078125, or 0 µL. Each well contained either one or two standard curves. A TissueLyser II was used to rupture tissues and plates were centrifuged for 2 minutes at 2000 rpm. 20µl of lysate from each well was added to 180µl of 1xPBS in a Black/Clear Bottom Plate 96-well plate (Thermo Scientific 265301). A spectrophotometer was used to measure the fluorescence intensity in each well using the 485/520 emission channel. Standard curves were used to calculate the sucrose concentration in each well. In our unfed controls we calculated sucrose readings at a maximum of 0.05µl so we used 0.06µl as the minimum value for considering an animal as fed.

Bugdorm BD4M1515) with access to 10% sucrose mixed with fluorescein at a final concentration of 0.002%. In experiments with capsaicin, mosquitoes were offered either a control meal consisting of 10% sucrose, 0.002% fluorescein and 0.01% DMSO (which was used as a vehicle to for capsaicin) or a meal consisting of 10% sucrose, 0.002% fluorescein and 50µM capsaicin (Tocris Bioscience - 0462/100). 32 females were not offered a fluorescein meal and reserved as unfed controls. Mosquitoes were allowed to feed for 4 hours. At the end of each trial mosquitoes were frozen at −20 °C until meal quantification was done.

### Single choice meal size quantification

Meal size quantification was done according to (Venkataraman et al., 2022). Animals that were offered a fluorescein meal as well as 8-16 unfed controls were added to a 96-well plate with one 2mm diameter stainless steel bead and 100µl of 1x PBS. 8 animals that were not offered a meal were used to generate a standard curve by adding the following volumes of 10% sucrose + 0.002% fluorescein: 5, 2.5, 1.25, 0.625, 0.3125, 0.15625, 0.078125, or 0 µL. Each well contained either one or two standard curves. A TissueLyser II was used to rupture tissues and plates were centrifuged for 2 minutes at 2000 rpm. 20µl of lysate from each well was added to 180µl of 1xPBS in a Black/Clear Bottom Plate 96-well plate (Thermo Scientific 265301). A spectrophotometer was used to measure the fluorescence intensity in each well using the 485/520 emission channel. Standard curves were used to calculate the sucrose concentration in each well. In our unfed controls we calculated sucrose readings at a maximum of 0.05µl so we used 0.06µl as the minimum value for considering an animal as fed.

**Figure S1.**
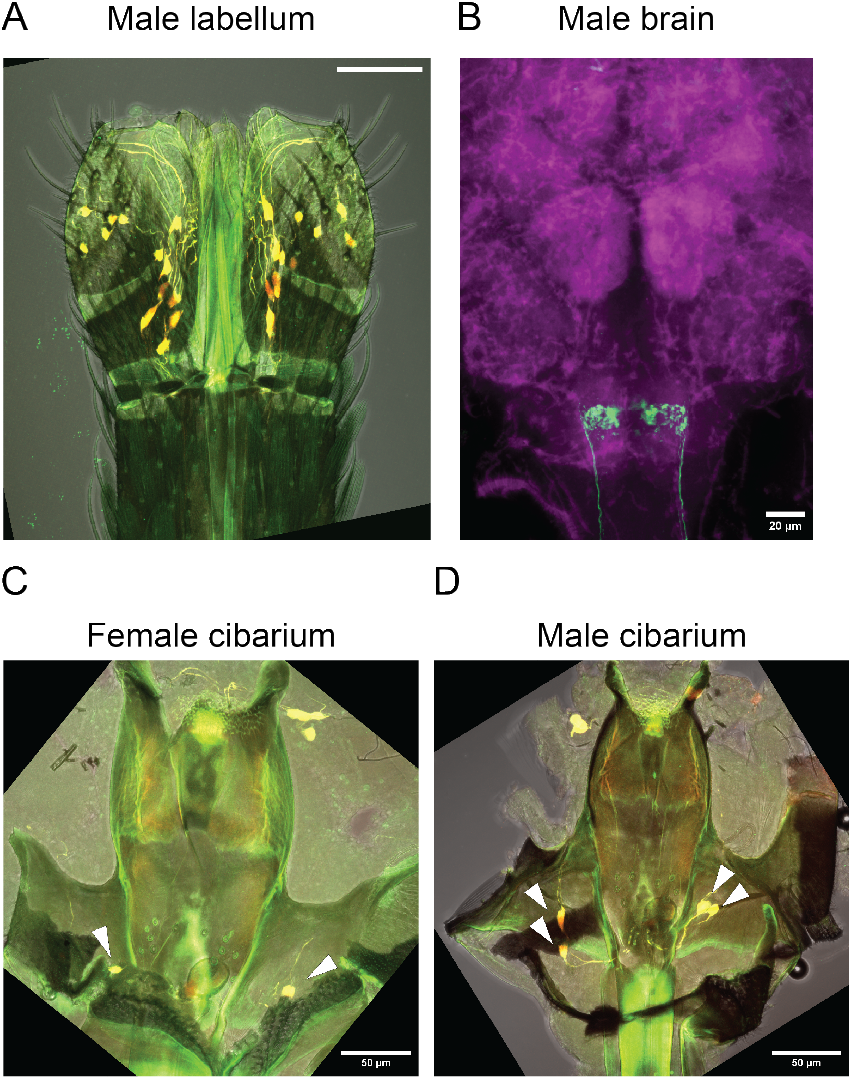
expression of Gr14 in male labellum and male and female cibarium. **(A)** Image of male labellum and labrum from *Aaeg*^*QF2*^ crossed to *QUAS-dTomato-T2A-GCaMP6s* (Matthews et al., 2019) shows gustatory receptor neurons (GRNs) labelled with QF2-driven fluorescence. **(E)** Immunohisto-chemical localization of *Gr14*-expressing GRN axons in the subesophogeal zone (SEZ) of the male brain. **(C)** Localization of *Gr14*-expressing neurons in dissected cibariums from female and **(D)** male mosquitoes. Scale bars: 50µm (A, C-D); 20µm B.

**Figure S2.**
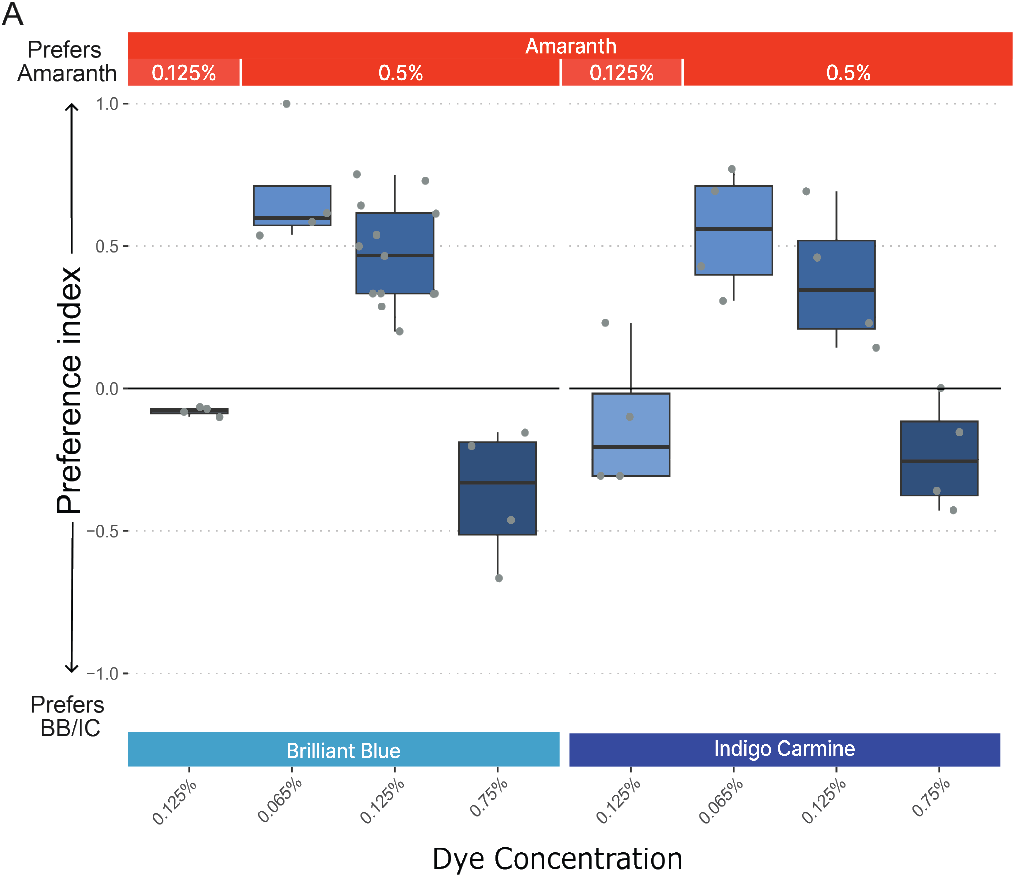
Mosquitoes show neutral preference for dyes at low concentrations. **(A)** The feeding preferences of female mosquitoes offered a sucrose meal with either Amaranth at varying concentrations, or Brilliant Blue (BB) dye or Indigo Carmine (IC) at varying concentrations. Feeding preference is indicated by a preference index, where 1.0 = complete preference for Amaranth dye and −1.0 = a complete preference for either Brilliant Blue dye or Indigo Carmine. Boxplots are showing the median and upper and lower quartiles. A mix of wild-type (*+/+*) and heterozygous (*Gr14*^*QF2/+*^) mosquitoes were used in this experiment.

**Figure S3.**
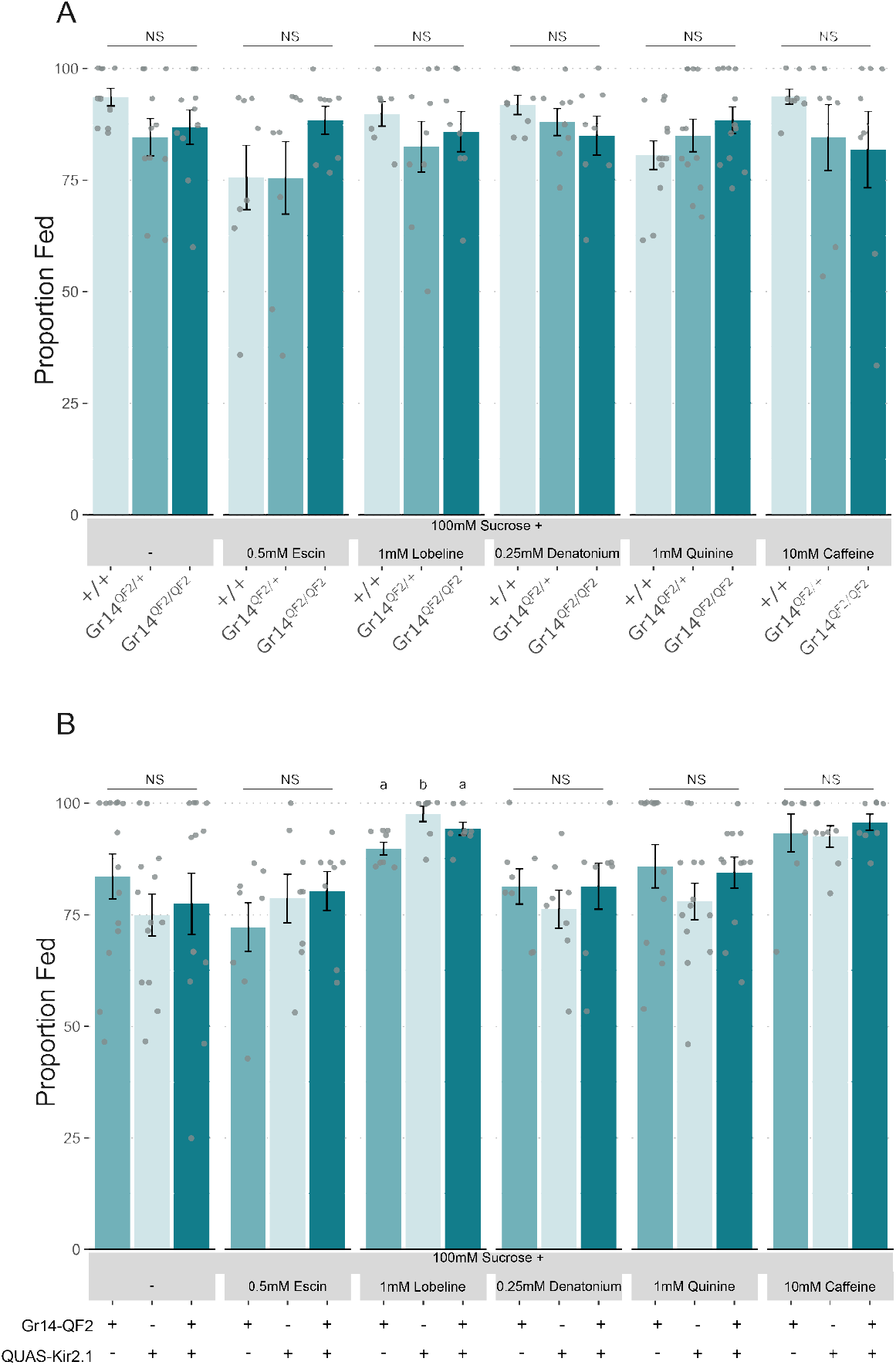
Knocking out *Gr14* or silencing *Gr14* neurons does not affect feeding rates when bitters are added to a nectar meal. **(A)** Proportion of mosquitoes that fed in the mutant nectar-feeding assay for wildtype (+/+) heterozygotes (*Gr14*^*QF2/+*^) and homoallelic Gr14 mutants (*Gr14*^*QF2/QF2*^) across the indicated bitter treatments. **(B)** Proportion of mosquitoes that fed in the sugar-feeding assay for driver controls (*Gr14-QF2*), reporter controls (*QUAS-Kir2*.*1*), and Gr14-silenced mosquitoes (*Gr14 > Kir2*.*1*). Points represent individual cages and bars represent mean ± SEM (n = 8 trials of 9–12 mosquitoes per cage). Letters above bars indicate significant differences between genotypes within each treatment, while NS indicates no significant difference (Kruskal–Wallis test followed by a Dunn’s test with a Bonferroni adjustment for multiple comparisons, P < 0.05).

**Figure S4.**
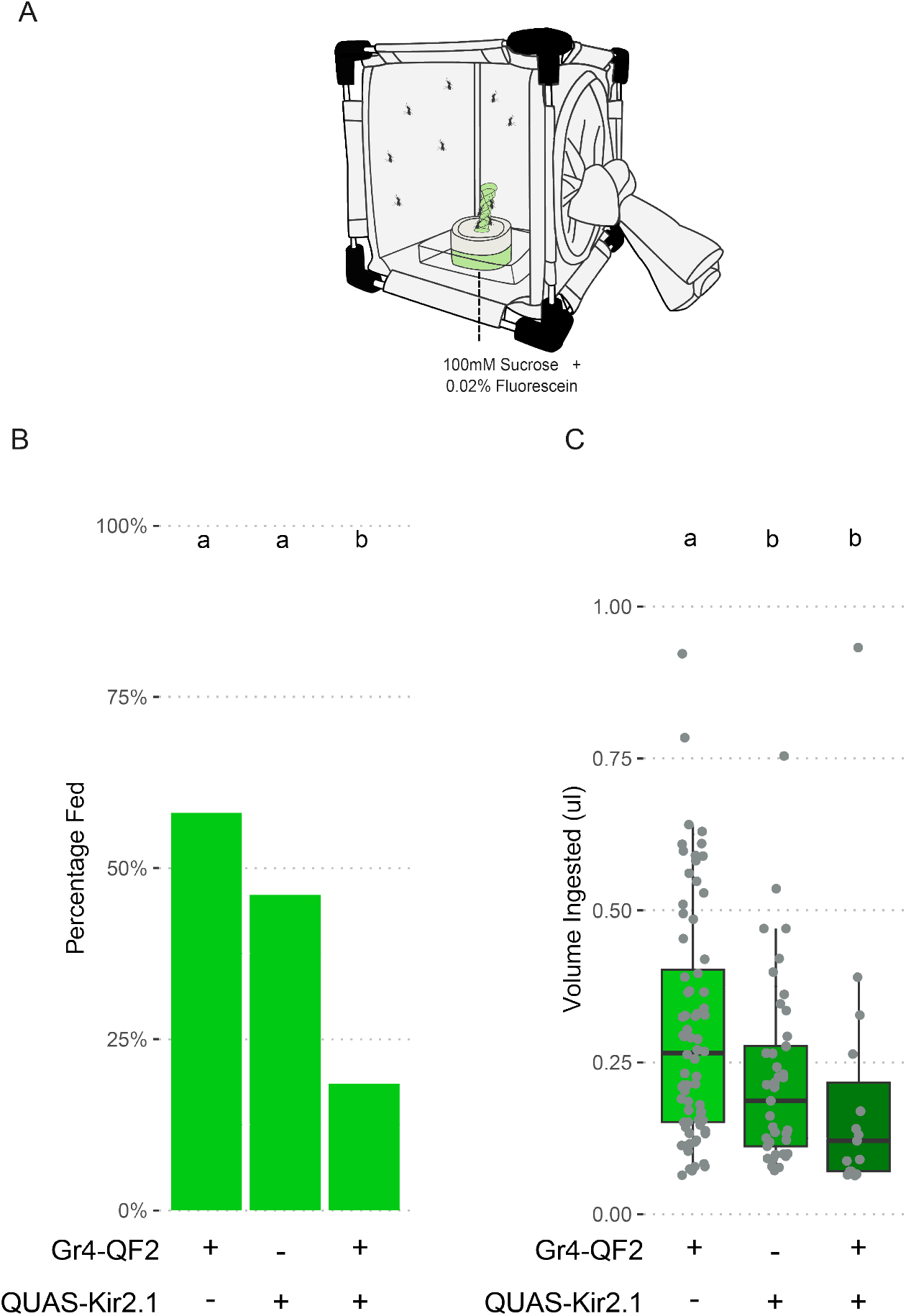
Silencing sugar neurons using Kir2.1 reduces sucrose ingestion. **(A)** Schematic of the single choice feeding assay where mosquitoes are offered a solution of 100mM sucrose, and 0.02% fluorescein **(B)** Percentage of mosquitoes that ingested 10% sucrose in 4 hours after a 24-hour water starvation. The genotypes included are driver controls (*Gr4-QF2*, n = 169), reporter controls (*QUAS-Kir2*.*1*, n = 116) and mosquitoes where Gr4 neurons are silenced (*Gr4 > Kir2*.*1*, n =111). **(B)** The volume of sucrose ingested by mosquitoes scored as fed. Letters indicate significant differences between genotypes: in **(A)** logistic regression followed by Tukey HSD, in **(B)** Kruskall-Wallis test followed by Dunn’s test with a Bonferroni adjustment for multiple comparisons (*P* < 0.05).

**Figure S5.**
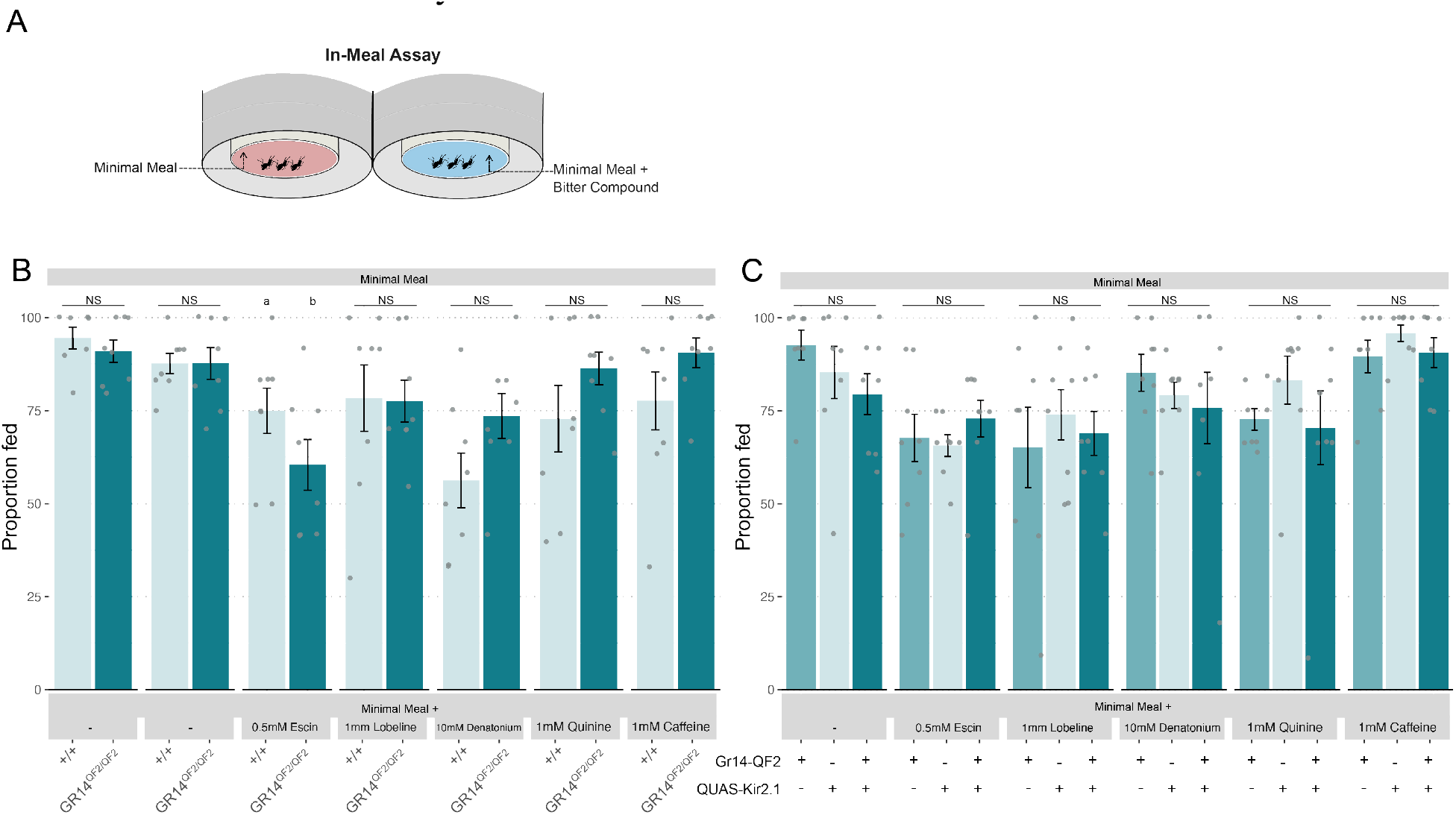
Knocking out *Gr14* or silencing *Gr14* expressing neurons does not alter feeding proportion when bitters are added directly to a bloodmeal. **(A)** Schematic of the In-Meal feeding assay, in which mosquitoes choose between a minimal meal and a minimal meal supplemented with a bitter compound. **(B)** Proportion of mosquitoes that fed in the In-Meal assay for wildtype (+/+) and homoallelic Gr14 mutants (*Gr14*^*QF2/QF2*^) across the indicated bitter treatments. **(C)** Proportion of mosquitoes that fed in the In-Meal assay for driver controls (*Gr14-QF2*), reporter controls (*QUAS-Kir2*.*1*) and mosquitoes where *Gr14* expressing neurons are silenced (*Gr14 > Kir2*.*1*).Points represent individual cages and bars represent mean ± SEM (n = 8 trials of 9–12 mosquitoes per cage). Letters above bars indicate significant differences between genotypes within each treatment, while NS indicates no significant difference (Kruskal– Wallis test followed by Dunn’s multiple comparisons test, P < 0.05).

**Figure S6.**
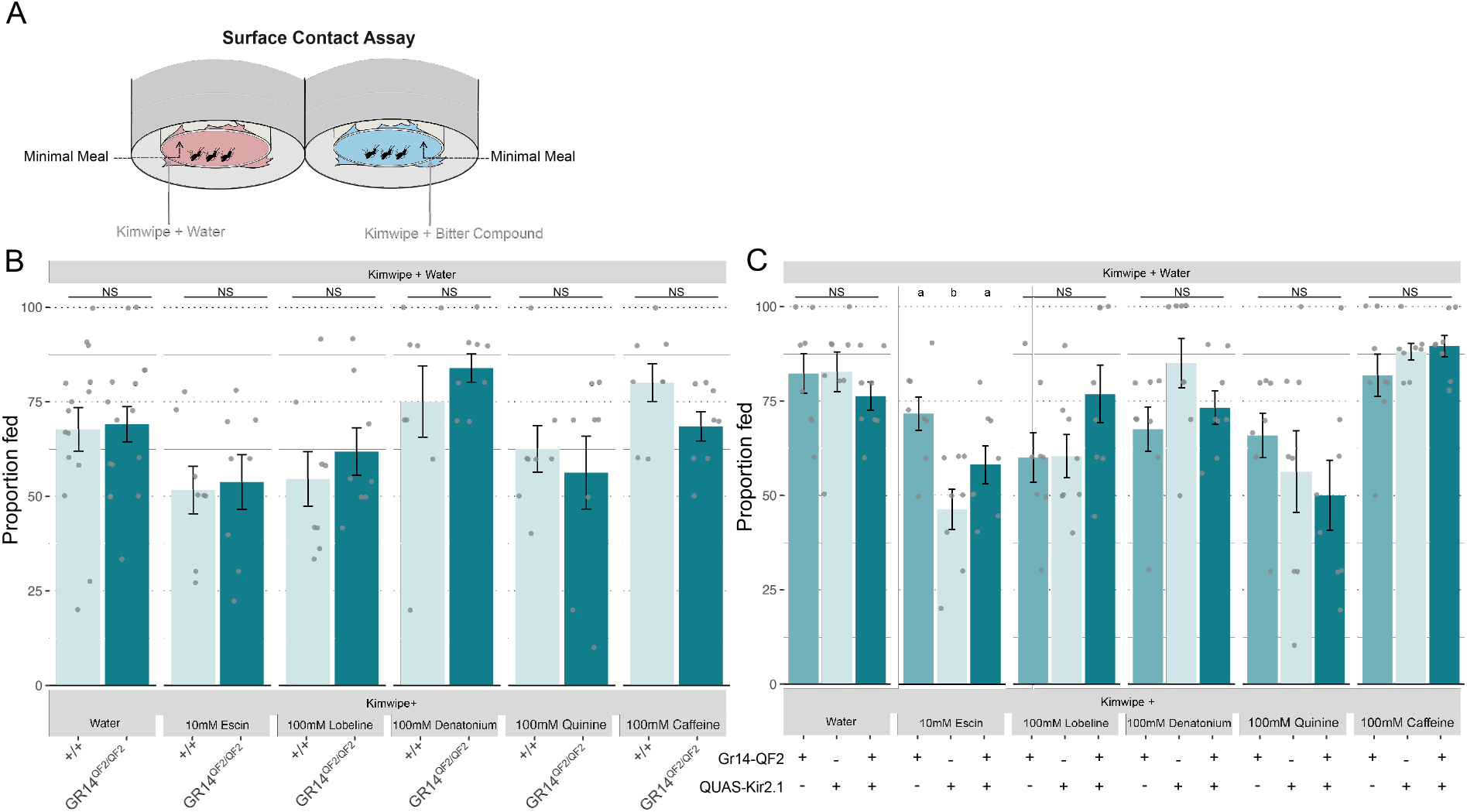
Knocking out *Gr14* or silencing *Gr14* expressing neurons does not alter feeding proportion when bitters are added to the surface of a bloodmeal. **(A)** Schematic of the Surface Contact assay, in which mosquitoes contact a Kimwipe treated with water or an indicated bitter compound prior to feeding on control minimal meals. **(B)** Proportion of mosquitoes that fed in the Surface Contact assay for wildtype (+/+) and homoallelic Gr14 mutants (*Gr14*^*QF2/QF2*^). **(C)** Proportion of mosquitoes that fed in the Surface Contact assay for driver controls (*Gr14-QF2*), reporter controls (*QUAS-Kir2*.*1*) and mosquitoes where *Gr14* expressing neurons are silenced (*Gr14 > Kir2*.*1*). Points represent individual cages and bars represent mean ± SEM (n = 8 trials of 9–12 mosquitoes per cage). Letters above bars indicate significant differences between genotypes within each treatment, while NS indicates no significant difference (Kruskal– Wallis test followed by Dunn’s multiple comparisons test, P < 0.05).

## Notes

### Competing Interest Statement

The authors have declared no competing interest.

## References

Adler, L.S., 2000. The ecological significance of toxic nectar. Oikos 91, 409–420. 10.1034/j.1600-0706.2000.910301.x

Baik, L.S., Carlson, J.R., 2020. The mosquito taste system and disease control. Proc. Natl. Acad. Sci. 117, 32848–32856. 10.1073/pnas.2013076117

Baik, L.S., Talross, G.J.S., Gray, S., Pattisam, H.S., Peterson, T.N., Nidetz, J.E., Hol, F.J.H., Carlson, J.R., 2024. Mosquito taste responses to human and floral cues guide biting and feeding. Nature 635, 639–646. 10.1038/s41586-024-08047-y

Baker, H.G., 1977. Non-sugar chemical constituents of nectar. Apidologie 8, 349–356. 10.1051/apido:19770405

Barredo, E., DeGennaro, M., 2020. Not Just from Blood: Mosquito Nutrient Acquisition from Nectar Sources. Trends Parasitol. 36, 473–484. 10.1016/j.pt.2020.02.003

Bernays, E.A., 1998. Evolution of Feeding Behavior in Insect Herbivores. BioScience 48, 35–44. 10.2307/1313226

Biere, A., Marak, H.B., van Damme, J.M.M., 2004. Plant chemical defense against herbivores and pathogens: generalized defense or trade-offs? Oecologia 140, 430–441. 10.1007/s00442-004-1603-6

Clements, A.N., 1992. The Biology of Mosquitoes: Vol. 1 Development, Nutrition and Reproduction. CABI.

Clyne, P.J., Warr, C.G., Carlson, J.R., 2000. Candidate taste receptors in Drosophila. Science 287, 1830–1834. 10.1126/science.287.5459.1830

Corfas, R.A., Vosshall, L.B., 2015. The cation channel TRPA1 tunes mosquito thermotaxis to host temperatures. eLife 4, e11750. 10.7554/eLife.11750

Coutinho-Abreu, I.V., Riffell, J.A., Akbari, O.S., 2022. Human attractive cues and mosquito host-seeking behavior. Trends Parasitol. 38, 246–264. 10.1016/j.pt.2021.09.012

DeGennaro, M., McBride, C.S., Seeholzer, L., Nakagawa, T., Dennis, E.J., Goldman, C., Jasinskiene, N., James, A.A., Vosshall, L.B., 2013. orco mutant mosquitoes lose strong preference for humans and are not repelled by volatile DEET. Nature 498, 487–491. 10.1038/nature12206

Delventhal, R., Carlson, J.R., 2016. Bitter taste receptors confer diverse functions to neurons. eLife 5, e11181. 10.7554/eLife.11181

Dennis, E.J., Goldman, O.V., Vosshall, L.B., 2019. Aedes aegypti Mosquitoes Use Their Legs to Sense DEET on Contact. Curr. Biol. 29, 1551–1556.e5. 10.1016/j.cub.2019.04.004

Dweck, H.K.M., Carlson, J.R., 2020. Molecular Logic and Evolution of Bitter Taste in Drosophila. Curr. Biol. 30, 17–30.e3. 10.1016/j.cub.2019.11.005

Foster, W.A., 1995. Mosquito sugar feeding and reproductive energetics. Annu. Rev. Entomol. 40, 443–474. 10.1146/annurev.en.40.010195.002303

French, A.S., Sellier, M.-J., Ali Agha, M., Guigue, A., Chabaud, M.-A., Reeb, P.D., Mitra, A., Grau, Y., Soustelle, L., MarionPoll, F., 2015. Dual Mechanism for Bitter Avoidance in Drosophila. J. Neurosci. 35, 3990–4004. 10.1523/JNEUROSCI.1312-14.2015

Glendinning, J.I., 1994. Is the bitter rejection response always adaptive? Physiol. Behav. 56, 1217–1227. 10.1016/0031-9384(94)90369-7

Goldman, O.V., DeFoe, A.E., Qi, Y., Jiao, Y., Weng, S.-C., Wick, B., Houri-Zeevi, L., Lakhiani, P., Morita, T., Razzauti, J., Rosas-Villegas, A., Tsitohay, Y.N., Walker, M.M., Hopkins, B.R., Ang, J.X.D., Antoshechkin, I., Cai, Y., Chen, F., Chen, Y.-C., Devilliers, J., Dong, L., Feuda, R., Gabrieli, P., Kopp, A., Kwon, H., Li, H.-H., Lu, T.-C., Lucio, T., Marques, J.T., Oliveira, M.F., Olmo, R.P., Palatini, U., Pithawala, Z.M., Pompon, J., Reis, Y., Rodrigues, J., Smith, R.C., Haeussler, M., Akbari, O.S., Duvall, L.B., White-Cooper, H., Sorrells, T.R., Sharma, R., Li, H., Vosshall, L.B., Shai, N., 2025. A single-nucleus transcriptomic atlas of the adult Aedes aegypti mosquito. Cell S0092867425011377. 10.1016/j.cell.2025.10.008

Gomes, J.V., Singh-Bhagania, S., Cenci, M., Chacon Cordon, C., Singh, M., Butterwick, J.A., 2024. The molecular basis of sugar detection by an insect taste receptor. Nature 629, 228–234. 10.1038/s41586-024-07255-w

Harrell, R.A., 2024. Mosquito Embryo Microinjection under Halo-carbon Oil or in Aqueous Solution. Cold Spring Harb. Protoc. 2024, pdb.prot108203. 10.1101/pdb.prot108203

Herre, M., Goldman, O.V., Lu, T.-C., Caballero-Vidal, G., Qi, Y., Gilbert, Z.N., Gong, Z., Morita, T., Rahiel, S., Ghaninia, M., 2022. Non-canonical odor coding in the mosquito. Cell 185, 3104–3123.

Ibanez, S., Gallet, C., Després, L., 2012. Plant Insecticidal Toxins in Ecological Networks. Toxins 4, 228–243. 10.3390/toxins4040228

Ignell, R., Hansson, B.S., 2005. Projection patterns of gustatory neurons in the suboesophageal ganglion and tritocerebrum of mosquitoes. J. Comp. Neurol. 492, 214–233. 10.1002/cne.20691

Ignell, R., Okawa, S., Englund, J.-E., Hill, S.R., 2010. Assessment of diet choice by the yellow fever mosquito Aedes aegypti. Physiol. Entomol. 35, 274–286. 10.1111/j.1365-3032.2010.00740.x

Jeong, Y.T., Shim, J., Oh, S.R., Yoon, H.I., Kim, C.H., Moon, S.J., Montell, C., 2013. An odorant-binding protein required for suppression of sweet taste by bitter chemicals. Neuron 79, 725–737. 10.1016/j.neuron.2013.06.025

Johnson, S.D., Hargreaves, A.L., Brown, M., 2006. Dark, bitter-tasting nectar functions as a filter of flower visitors in a bird-pollinated plant. Ecology 87, 2709–2716. 10.1890/0012-9658(2006)87%5B2709:dbnfaa%5D2.0.co;2

Jové, V., Gong, Z., Hol, F.J.H., Zhao, Z., Sorrells, T.R., Carroll, T.S., Prakash, M., McBride, C.S., Vosshall, L.B., 2020a. Sensory Discrimination of Blood and Floral Nectar by Aedes aegypti Mosquitoes. Neuron 108, 1163–1180.e12. 10.1016/j.neuron.2020.09.019

Jové, V., Venkataraman, K., Gabel, T.M., Duvall, L.B., 2020b. Feeding and Quantifying Animal-Derived Blood and Artificial Meals in Aedes aegypti Mosquitoes. J. Vis. Exp. 61835. 10.3791/61835

Kent, L.B., Walden, K.K.O., Robertson, H.M., 2008. The Gr Family of Candidate Gustatory and Olfactory Receptors in the Yellow-Fever Mosquito Aedes aegypti. Chem. Senses 33, 79–93. 10.1093/chemse/bjm067

Kessler, S., Vlimant, M., Guerin, P.M., 2013. The sugar meal of the African malaria mosquito Anopheles gambiae and how deterrent compounds interfere with it: a behavioural and neurophysiological study. J. Exp. Biol. 216, 1292–1306. 10.1242/jeb.076588

Kistler, K.E., Vosshall, L.B., Matthews, B.J., 2015. Genome Engineering with CRISPR-Cas9 in the Mosquito Aedes aegypti. Cell Rep. 11, 51–60. 10.1016/J.CELREP.2015.03.009

Kwon, Y., Kim, S.H., Ronderos, D.S., Lee, Y., Akitake, B., Wood-ward, O.M., Guggino, W.B., Smith, D.P., Montell, C., 2010. Drosophila TRPA1 Channel Is Required to Avoid the Naturally Occurring Insect Repellent Citronellal. Curr. Biol. 20, 1672–1678. 10.1016/j.cub.2010.08.016

Laursen, W.J., 2026. The sensory biology of mosquito gustation. Curr. Opin. Neurobiol. 99, 103215. 10.1016/j.conb.2026.103215

Lazzari, C.R., Ortega-Insaurralde, I., Esnault, J., Costa, E., Crespo, J.E., Barrozo, R.B., 2024. Mosquitoes do not Like Bitter. J. Chem. Ecol. 50, 143–151. 10.1007/s10886-024-01476-z

Lee, Y., Moon, S.J., Montell, C., 2009. Multiple gustatory receptors required for the caffeine response in Drosophila. Proc. Natl. Acad. Sci. 106, 4495–4500. 10.1073/pnas.0811744106

Li, M., Bui, M., Yang, T., Bowman, C.S., White, B.J., Akbari, O.S., 2017. Germline Cas9 expression yields highly efficient genome engineering in a major worldwide disease vector, Aedes aegypti. Proc. Natl. Acad. Sci. U. S. A. 114, E10540–E10549. 10.1073/pnas.1711538114

Lüttge, U., 1977. NECTAR COMPOSITION AND MEMBRANE TRANSPORT OF SUGARS AND AMINO ACIDS: A REVIEW ON THE PRESENT STATE OF NECTAR RESEARCH. Apidologie 8, 305–319. 10.1051/apido:19770402

Ma, D., Hu, M., Yang, X., Liu, Q., Ye, F., Cai, W., Wang, Y., Xu, X., Chang, S., Wang, R., Yang, W., Ye, S., Su, N., Fan, M., Xu, H., Guo, J., 2024. Structural basis for sugar perception by Drosophila gustatory receptors. Science 383, eadj2609. 10.1126/science.adj2609

Marella, S., Fischler, W., Kong, P., Asgarian, S., Rueckert, E., Scott, K., 2006. Imaging taste responses in the fly brain reveals a functional map of taste category and behavior. Neuron 49, 285–295. 10.1016/j.neuron.2005.11.037

Matthews, B.J., Dudchenko, O., Kingan, S.B., Koren, S., Antoshechkin, I., Crawford, J.E., Glassford, W.J., Herre, M., Redmond, S.N., Rose, N.H., Weedall, G.D., Wu, Y., Batra, S.S., Brito-Sierra, C.A., Buckingham, S.D., Campbell, C.L., Chan, S., Cox, E., Evans, B.R., Fansiri, T., Filipović, I., Fontaine, A., Gloria-Soria, A., Hall, R., Joardar, V.S., Jones, A.K., Kay, R.G.G., Kodali, V.K., Lee, J., Lycett, G.J., Mitchell, S.N., Muehling, J., Murphy, M.R., Omer, A.D., Partridge, F.A., Peluso, P., Aiden, A.P., Ramasamy, V., Rašić, G., Roy, S., Saavedra-Rodriguez, K., Sharan, S., Sharma, A., Smith, M.L., Turner, J., Weakley, A.M., Zhao, Z., Akbari, O.S., Black, W.C., Cao, H., Darby, A.C., Hill, C.A., Johnston, J.S., Murphy, T.D., Raikhel, A.S., Sattelle, D.B., Sharakhov, I.V., White, B.J., Zhao, L., Aiden, E.L., Mann, R.S., Lambrechts, L., Powell, J.R., Sharakhova, M.V., Tu, Z., Robertson, H.M., McBride, C.S., Hastie, A.R., Korlach, J., Neafsey, D.E., Phillippy, A.M., Vosshall, L.B., 2018. Improved reference genome of Aedes aegypti informs arbovirus vector control. Nature 563, 501–507. 10.1038/s41586-018-0692-z

Matthews, B.J., McBride, C.S., DeGennaro, M., Despo, O., Vosshall, L.B., 2016. The neurotranscriptome of the Aedes aegypti mosquito. BMC Genomics 17, 32. 10.1186/s12864-015-2239-0

Matthews, B.J., Younger, M.A., Vosshall, L.B., 2019. The ion channel ppk301 controls freshwater egg-laying in the mosquito aedes aegypti. eLife 8. 10.7554/eLife.43963

McIver, S., Siemicki, R., 1978. Fine structure of tarsal sensilla ofAedes aegypti (L.) (Diptera: Culicidae). J. Morphol. 155, 137–155. 10.1002/jmor.1051550202

Mclver, S.B., 1982. Sensilla of Mosquitoes (Diptera: Culicidae). J. Med. Entomol. 19, 489–535. 10.1093/jmedent/19.5.489

McMeniman, C.J., Corfas, R.A., Matthews, B.J., Ritchie, S.A., Vosshall, L.B., 2014. Multimodal integration of carbon dioxide and other sensory cues drives mosquito attraction to humans. Cell 156, 1060–1071. 10.1016/j.cell.2013.12.044

Melo, N., Capek, M., Arenas, O.M., Afify, A., Yilmaz, A., Potter, C.J., Laminette, P.J., Para, A., Gallio, M., Stensmyr, M.C., 2021. The irritant receptor TRPA1 mediates the mosquito repellent effect of catnip. Curr. Biol. 31, 1988–1994.e5. 10.1016/j.cub.2021.02.010

Montell, C., 2025. The sensory arsenal mosquitoes use to find us. Trends Parasitol. 41, 591–602. 10.1016/j.pt.2025.05.004

Moon, S.J., Köttgen, M., Jiao, Y., Xu, H., Montell, C., 2006. A Taste Receptor Required for the Caffeine Response In Vivo. Curr. Biol. 16, 1812–1817. 10.1016/j.cub.2006.07.024

Muñoz, I.J., Schilman, P.E., Barrozo, R.B., 2020. Impact of alkaloids in food consumption, metabolism and survival in a blood-sucking insect. Sci. Rep. 10, 9443. 10.1038/s41598-020-65932-y

Owen, W.B., Larsen, J.R., Pappas, L.G., 1974. Functional units in the labellar chemosensory hairs of the mosquito Culiseta inornata (Williston). J. Exp. Zool. 188, 235–247. 10.1002/jez.1401880211

Pappas, L.G., Larsen, J.R., 1976. Gustatory hairs on the mosquito, Culiseta inornata. J. Exp. Zool. 196, 351–360. 10.1002/jez.1401960309

Peach, D.A.H., Gries, G., 2020. Mosquito phytophagy – sources exploited, ecological function, and evolutionary transition to haematophagy. Entomol. Exp. Appl. 168, 120–136. 10.1111/eea.12852

Riabinina, O., Task, D., Marr, E., Lin, C.-C., Alford, R., O’Brochta, D.A., Potter, C.J., 2016. Organization of olfactory centres in the malaria mosquito Anopheles gambiae. Nat. Commun. 7, 13010. 10.1038/ncomms13010

Robertson, H.M., 2019. Molecular evolution of the major arthropod chemoreceptor gene families. Annu. Rev. Entomol. 64, 227–242. 10.1146/annurev-ento-020117-043322

Robertson, H.M., Warr, C.G., Carlson, J.R., 2003. Molecular evolution of the insect chemoreceptor gene superfamily in Drosophila melanogaster. Proc. Natl. Acad. Sci. 100, 14537–14542. 10.1073/pnas.2335847100

Sanford, J.L., Shields, V.D.C., Dickens, J.C., 2013. Gustatory receptor neuron responds to DEET and other insect repellents in the yellow-fever mosquito, Aedes aegypti. Naturwissenschaften 100, 269–273. 10.1007/s00114-013-1021-x

Shannon, D.M., Richardson, N., Lahondère, C., Peach, D., 2024. Mosquito floral visitation and pollination. Curr. Opin. Insect Sci. 65, 101230. 10.1016/j.cois.2024.101230

Shim, J., Lee, Y., Jeong, Y.T., Kim, Y., Lee, M.G., Montell, C., Moon, S.J., 2015. The full repertoire of Drosophila gustatory receptors for detecting an aversive compound. Nat. Commun. 6, 8867. 10.1038/ncomms9867

Sparks, J.T., Dickens, J.C., 2016a. Bitter-sensitive gustatory receptor neuron responds to chemically diverse insect repellents in the common malaria mosquito Anopheles quadrimaculatus. Naturwissenschaften 103, 39. 10.1007/s00114-016-1367-y

Sparks, J.T., Dickens, J.C., 2016b. Electrophysiological Responses of Gustatory Receptor Neurons on the Labella of the Common Malaria Mosquito, Anopheles quadrimaculatus (Diptera: Culicidae). J. Med. Entomol. 53, 1148–1155. 10.1093/jme/tjw073

Sparks, J.T., Vinyard, B.T., Dickens, J.C., 2013. Gustatory receptor expression in the labella and tarsi of Aedes aegypti. Insect Biochem. Mol. Biol. 43, 1161–1171. 10.1016/j.ibmb.2013.10.005

Stern, D.L., 2016. Tagmentation-Based Mapping (TagMap) of Mobile DNA Genomic Insertion Sites. bioRxiv. 10.1101/037762

Sung, H.Y., Jeong, Y.T., Lim, J.Y., Kim, H., Oh, S.M., Hwang, S.W., Kwon, J.Y., Moon, S.J., 2017. Heterogeneity in the Drosophila gustatory receptor complexes that detect aversive compounds. Nat. Commun. 8, 1484. 10.1038/s41467-017-01639-5

Thorne, N., Chromey, C., Bray, S., Amrein, H., 2004. Taste Perception and Coding in Drosophila. Curr. Biol. 14, 1065–1079. 10.1016/j.cub.2004.05.019

Venkataraman, K., Jové, V., Duvall, L.B., 2022. Size Quantification of Blood and Sugar Meals in Aedes aegypti Mosquitoes. Cold Spring Harb. Protoc. 2022, pdb.prot107862. 10.1101/pdb.prot107862

Wang, Z., Singhvi, A., Kong, P., Scott, K., 2004. Taste Representations in the Drosophila Brain. Cell 117, 981–991. 10.1016/j.cell.2004.06.011

Weiss, L.A., Dahanukar, A., Kwon, J.Y., Banerjee, D., Carlson, J.R., 2011. The molecular and cellular basis of bitter taste in Drosophila. Neuron 69, 258–272. 10.1016/j.neuron.2011.01.001

Yang, C.-H., Rumpf, S., Xiang, Y., Gordon, M.D., Song, W., Jan, L.Y., Jan, Y.-N., 2009. Control of the postmating behavioral switch in Drosophila females by internal sensory neurons. Neuron 61, 519–526. 10.1016/j.neuron.2008.12.021

Yarmolinsky, D.A., Zuker, C.S., Ryba, N.J.P., 2009. Common sense about taste: from mammals to insects. Cell 139, 234–244. 10.1016/j.cell.2009.10.001

Younger, M.A., 2024. Whole-Mount Immunofluorescent Labeling of the Mosquito Central Nervous System. Cold Spring Harb. Protoc. 2024, db.prot108336. 10.1101/pdb.prot108336

Ziaikin, E., David, M., Uspenskaya, S., Niv, M.Y., 2025. BitterDB: 2024 update on bitter ligands and taste receptors. Nucleic Acids Res. 53, D1645–D1650. 10.1093/nar/gkae1044

